# HDAC6 and SSAT2 orchestrate acetyllysine metabolism and protein homeostasis

**DOI:** 10.64898/2026.05.22.727066

**Authors:** Fang Wu, Santhosh Paramasivam, Satish Noonepalle, Albert Batushansky, Miroslava Vosahlikova, Alejandro Villagra, Cyril Bařinka, Eyal Arbely

## Abstract

Lysine acetylation is a prevalent and dynamic posttranslational modification, yet the metabolic fate and physiological role of free *N* ^ε^-acetyl-L-lysine (AcK) remain unclear. Here, we identify opposing enzyme-catalyzed reactions linking lysine/AcK homeostasis to protein stability. We show that the first catalytic domain of HDAC6 stereospecifically deacetylates free AcK, establishing HDAC6 as an *N* ^ε^-acyl-lysine deacylase. AcK functions both as a recyclable lysine reservoir and as a signaling metabolite that elevates α-tubulin acetylation and suppresses cell migration. Conversely, SSAT2 acetylates free lysine in cells and undergoes substrate-dependent C-terminal autoacetylation. We demonstrate that C-terminal lysine acetylation constitutes a reversible proteasomal degradation signal that promotes SSAT2 degradation and may extend to additional proteins bearing a terminal lysine. Stabilization of SSAT2 by HDAC6-dependent deacetylation, together with intricate regulation affected by substrate availability, reveals a novel mechanism that couples amino acid recycling to cytoskeletal dynamics and proteostasis.

## INTRODUCTION

Across all domains of life, proteins are posttranslationally modified by the covalent attachment of a wide array of chemical groups to amino acid side chains. While much attention has been given to how posttranslational modifications (PTMs) regulate protein activity, localization, and turnover, equally fundamental questions remain underexplored: Can all modified amino acids be found as free metabolites? If so, what is their functional role and metabolic fate? For example, *N* ^ε^-lysine acetylation is a conserved PTM, central to transcriptional regulation, metabolism, and aging.^1–5^ Despite its ubiquity, the biochemical fate of *N* ^ε^-acetyl-L-lysine (AcK) as a free metabolite remains enigmatic, particularly in mammals. AcK has been detected in various organisms, including in human cells, serum, and urine, with evidence suggesting its dietary uptake and conversion to lysine.^6–16^ Given that lysine is an essential amino acid in animals and must be obtained from external sources, such as dietary proteins or the gut microbiome, the ability to salvage it from acetylated metabolites is an important component of amino acid homeostasis.

Protein acetylation is governed by the dynamic interplay between lysine acetyltransferases (KATs, ‘writers’) and lysine deacetylases (KDACs, ‘erasers’).^17^ While the human genome encodes more than 30 known KATs, only 18 KDACs have been identified to date, 11 of which are Zn^2+^-dependent deacetylases (HDAC1–11).^18,19^ Notably, although *N* ^ε^-acyl-lysine deacylase activity has been defined (EC 3.5.1.17),^20^ none of the known human KDACs have been shown to catalyze this reaction. Intriguingly, genome-wide association studies (GWAS) found an association between altered serum AcK levels and single-nucleotide polymorphisms (SNPs) within the coding regions of one KDAC, histone deacetylase 6 (HDAC6), and one KAT, spermidine/spermine-N1-acetyltransferase 2 (SSAT2; Figure S1A).^10,11,21,22^ The Zn^2+^-dependent deacetylase HDAC6 uniquely harbors two catalytic domains (CD1 and CD2; Figure S1B).^23,24^ While the deacetylase activity of HDAC6 is generally attributed to CD2,^23–29^ the physiological functions of CD1 remain unclear. That said, CD1 has been reported to selectively recognize C-terminal acetylated lysines.^23,29^ Human SSAT2 (hereafter SSAT2), a member of the GCN5-related *N*-acetyltransferases (GNAT) family, was identified by its sequence similarity to SSAT1.^30^ Unlike SSAT1, which acetylates polyamines, SSAT2 preferentially catalyzes the *N* ^ε^-acetylation of thialysine and, to a lesser extent, lysine.^31–34^ SSAT2 has also been implicated in the oxygen-dependent degradation of hypoxia-inducible factor 1α and the activation of NF-*κ*B.^35,36^

In this study, we analyzed reversible lysine acetylation from the perspective of the free modified amino acid and used chemical and biological approaches to define the cellular functions of AcK, establish the roles of HDAC6 and SSAT2 in maintaining AcK homeostasis, and uncover the regulatory mechanisms that govern the activities and stability of both enzymes.

## RESULTS

### HDAC6 CD1 catalyzes the deacetylation of AcK in vitro and in vivo

Guided by GWAS analysis (Figure S1A) and preliminary observations in our lab,^37^ we hypothesized that HDAC6 and SSAT2 form an opposing enzyme pair that mediates the interconversion between lysine and AcK (Figure 1A). The activity of SSAT2 as an acetyltransferase has been previously characterized,^31–34^ whereas the role of HDAC6 in free AcK hydrolysis has not been described in the literature. Given this lack of prior characterization, we began our investigation with HDAC6. Supporting our hypothesis, we found that incubation of AcK with human HDAC6 (hereafter HDAC6) results in AcK hydrolysis and release of acetate (Figures 1B, 1C, S1C, S1D, and S1E). To identify the catalytic domain in HDAC6 responsible for AcK hydrolysis, H216 or H611 were mutated to alanine, generating CD1- or CD2-inactive HDAC6, respectively (Figure S1B). Acetate formation measured following the addition of AcK to cleared lysates from HEK293T cells overexpressing HDAC6, or to immunoprecipitated (IPed) HDAC6 revealed that AcK is deacetylated by HDAC6 CD1, and not CD2 (Figures 1D, 1E, S1F, and S1G). Similar results were obtained in activity measurements of bacterially expressed and purified truncated versions (residues 84–835) of HDAC6 (Figures 1F, S1H, and S1I). Lastly, we measured the steady-state kinetics of recombinantly expressed and purified *Danio rerio* HDAC6 CD1 (zCD1) as a function of AcK concentration, and found a robust catalytic efficiency, with an apparent *k*_cat_/*K* _m_ of ∼1,900 M^-1^·sec^-1^ (Figure 1G). Based on these data, we conclude that HDAC6 CD1 deacetylates AcK in vitro.

**Figure 1.**
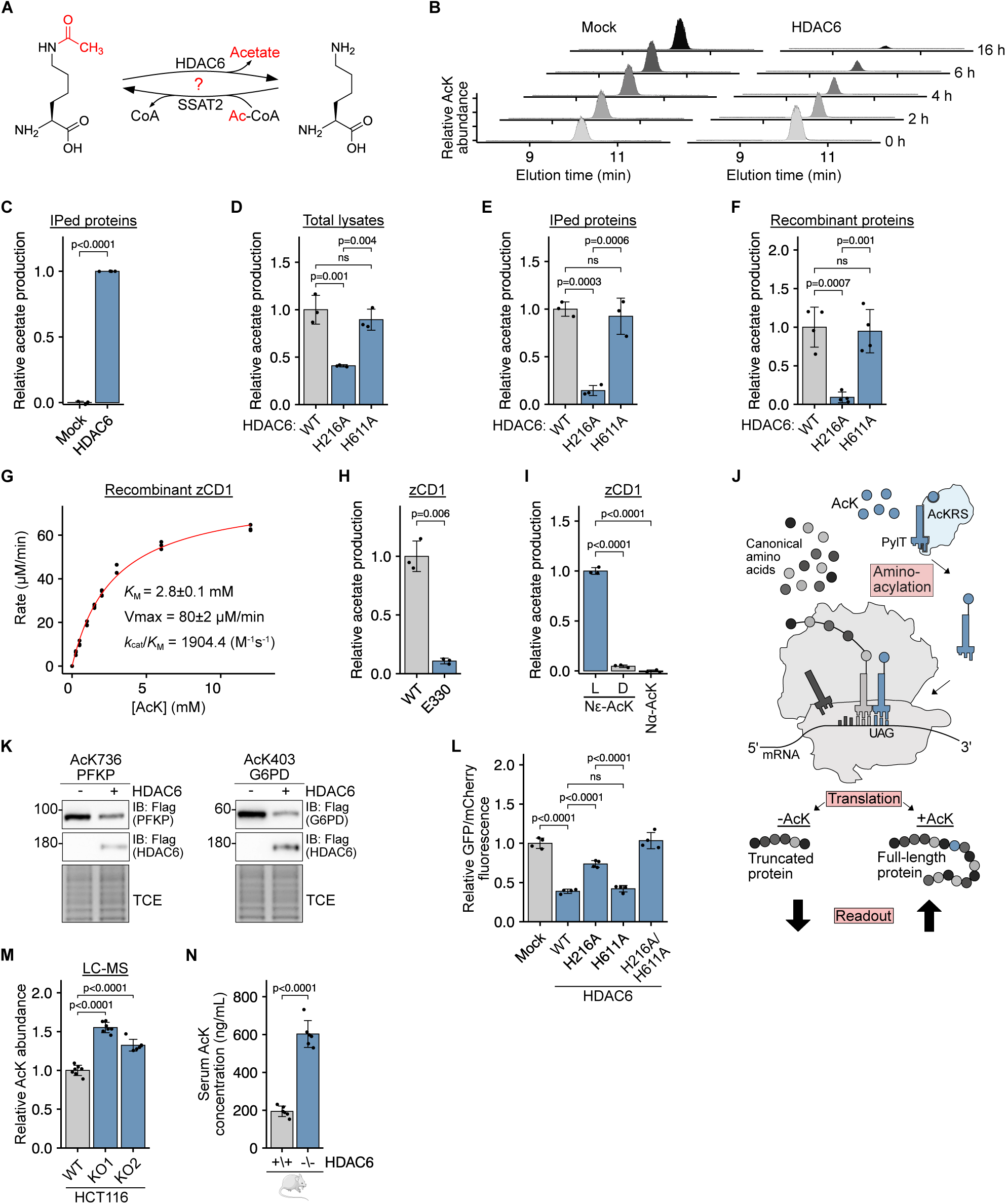
HDAC6 CD1 catalyzes the deacetylation of free *N* ^ε^-acetyl-L-lysine in vitro and in vivo. (A) AcK deacetylation and lysine acetylation reactions, suggested to be catalyzed by HDAC6 and SSAT2, respectively. (B) LC-MS analysis showing relative AcK abundance as a function of time, following incubation of AcK with Flag-isolated proteins from lysates of HEK293T cells transfected with HDAC6-Flag expressing plasmid or empty vector. (C) Relative acetate production following incubation of AcK with Flag-isolated proteins from lysates of HEK293T cells transfected with HDAC6-Flag expressing plasmid or empty vector. (D–F) Relative acetate production following incubation of AcK with WT or mutant HDAC6-Flag in cleared total lysates (D), with Flag-isolated WT or mutant HDAC6-Flag expressed in mammalian cells (E), or with purified WT or mutant hCD12 (aa 84–835) recombinantly expressed in *E. coli* (F). (G) Steady-state kinetics of AcK hydrolysis by purified zCD1. The red line represents the fitting of data to the Michaelis-Menten equation. (H) Relative acetate production following incubation of purified WT or E330 zCD1 with AcK. (I) Relative acetate production following incubation of purified zCD1 with *N ^ε^*-acetylated D- or L-lysine, or *N ^α^*-acetyl-L-lysine. (J) GCE-based assay for measuring relative cellular levels of AcK. Aminoacylation of PylT with AcK, catalyzed by an evolved AcKRS, enables the cotranslational incorporation of AcK and expression of a full-length reporter protein. (K) Immunoblotting using anti-Flag antibody, showing the levels of C-terminal Flag-tagged full-length proteins with an in-frame TAG mutation, co-expressed with AcKRS and with or without HDAC6-Flag. (L) Relative GFP/mCherry fluorescence ratio measured in lysates of HEK293T cells cultured with AcK and cotransfected with mCherry-TAG-GFP encoding plasmid and indicated HDAC6-Flag mutant or empty vector. (M) Relative abundance of AcK in WT and HDAC6-knockout HCT116 cell lines, measured by LC-MS. (N) Serum concentration of AcK in WT and HDAC6-knockout mice, measured by quantitative LC-MS. Data represent the mean ± S.D. of at least three independent experiments. Statistical analysis was performed using the Welch two-sample t-test (C, H, N) or ANOVA followed by Tukey post hoc analysis (D–F, I, L, M). ns, non-significant. In-gel TCE fluorescence serves as a loading control.

Having established AcK hydrolysis by HDAC6 CD1 in vitro, we next sought to understand the molecular basis for its substrate specificity. Previous studies found that in *Danio rerio* HDAC6 (zHDAC6), positively charged K330 (K353 in HDAC6) confers specificity for C-terminally acetylated lysine via an electrostatic interaction with the protein’s C-terminal carboxylate group.^23^ A structural model of AcK-bound zCD1 produced using Boltz-2, ^38^ suggests that AcK binding is facilitated by a similar interaction between K330 and the negatively charged carboxylate of AcK (Figure S1J). This was corroborated by inhibition of zCD1 activity towards AcK by the K330E mutation (Figure 1H). Moreover, significantly less acetate was formed when purified zCD1 was incubated with the D isomer of *N* ^ε^-acetyllysine, or *N* ^α^-acetyl-L-lysine, compared to the L isomer of *N* ^ε^-acetyllysine (Figure 1I). Consistently, HDAC6 expressed in mammalian cells also exhibited a higher preference for the L isomer (Figure S1K). Hence, HDAC6 CD1 catalyzes the deacetylation of *N* ^ε^-acetyl-L-lysine in a stereospecific manner.

To measure CD1 activity within the cellular environment, we established an assay for monitoring AcK levels in living cells (Figure 1J). The assay utilizes genetic code expansion (GCE) technology to co-translationally incorporate AcK in response to an in-frame UAG stop codon. In this system, the expression of a full-length protein encoded by a gene carrying an in-frame TAG mutation requires the aminoacylation of the amber-suppressor pyrrolysine tRNA (PylT) with AcK, by an evolved AcK synthetase (AcKRS). Thus, at subsaturating levels of AcK, normalized expression levels of a full-length protein (e.g., a fluorescent protein) serve as a semi-quantitative measure of AcK availability in the cell. Supporting our hypothesis, reduced full-length protein expression was observed in cells cultured with AcK when TAG-mutants of GFP and other proteins were coexpressed with HDAC6 (Figures 1K and S2A).

To enable better evaluation of AcK levels in living cells, we replaced GFP with a chimeric protein composed of mCherry and GFP, separated by an in-frame TAG stop codon (mCherry-TAG-GFP; Figure S2B). This dual-color assay enables a robust evaluation of relative cellular AcK levels in a multi-well format, as the GFP/mCherry fluorescence ratio correlates with cellular AcK availability and incorporation efficiency. Consistent with our observations, coexpression of HDAC6 with chimeric mCherry-TAG-GFP in cells cultured with AcK resulted in a significant decrease in GFP/mCherry fluorescence ratio (Figure S2C), due to lower GFP—rather than higher mCherry—expression levels (Figure S2D). The measured GFP/mCherry fluorescence ratio positively correlated with AcK concentration in the culture medium (Figure S2E), and inversely correlated with HDAC6 expression levels (Figures S2F). Importantly, inactivation of HDAC6 CD1, but not CD2, resulted in an increase in GFP/mCherry fluorescence ratio without affecting α-tubulin K40 acetylation, which is dependent on CD2 activity (Figures 1L, S2E, and S2G). These results validate our GCE-based assay and further confirm the activity of HDAC6 CD1 as AcK deacetylase in mammalian cells.

To determine the role of endogenous HDAC6 in AcK metabolism, we generated HDAC6-knockout HCT116 and HepG2 cell lines using clustered regularly interspaced short palindromic repeats (CRISPR)-Cas9 technology. For each cell line, two independent knockout lines were created using two different guide RNA sequences (Figures S2H and S2I). Liquid chromatography-mass spectrometry (LC-MS) analysis revealed significantly higher levels of AcK in HDAC6-knockout cell lines (Figures 1M, S2J, and S2K). Consistently, higher efficiency of AcK incorporation into GFP-N150TAG-HA was measured in HDAC6-knockout HCT116 cells (Figure S2L). We next sought to measure the effect of HDAC6 on AcK levels in vivo. Quantitative LC-MS measurements of AcK serum levels in WT and HDAC6-knockout mice found significantly higher concentrations of AcK in HDAC6-knockout mice compared to WT (600 ng/mL and 200 ng/mL, respectively), supporting a role for HDAC6 in systemic AcK homeostasis in mice (Figure 1N).

Taken together, these data provide new insight into the physiological role of the ambiguous CD1, and identify HDAC6 as a metabolic enzyme with *N* ^ε^-acyl-lysine deacylase activity (EC 3.5.1.17) attributed to its first catalytic domain in vitro and in cells, and implicate HDAC6 in systemic AcK homeostasis in vivo.

### AcK modulates CD2 activity and can serve as a lysine reservoir via recycling by CD1

While HDAC6 has been the subject of numerous studies, less is known about the metabolism and physiological roles of free acylated lysine derivatives in general, and AcK in particular. We first evaluated if representative KDACs from classes I–III are involved in AcK metabolism, and found that among the KDACs tested, only HDAC6 (class IIb) can catalyze the deacetylation of AcK in cells (Figures 2A). Next, we measured the ability of HDAC6 CD1 to catalyze the deacylation of other acylated lysine derivatives, as lysine residues can be modified by conjugation to different acyl groups. Using our dual-color assay with the most efficient synthetases, we found no significant CD1-dependent deacylation of propionyl lysine (PrK), 2-hydroxyisobutyryl lysine (HibK), and butyryl lysine (BuK), suggesting a higher preference of HDAC6 CD1 for AcK (Figure S3A, S3B, and S3C). The lack of effect on PrK incorporation further validates our GCE-based cellular assay since the incorporation of AcK and PrK is facilitated by the same evolved synthetase. Importantly, it highlights CD1’s involvement in AcK metabolism and its sensitivity to the identity of the acyl group.

**Figure 2.**
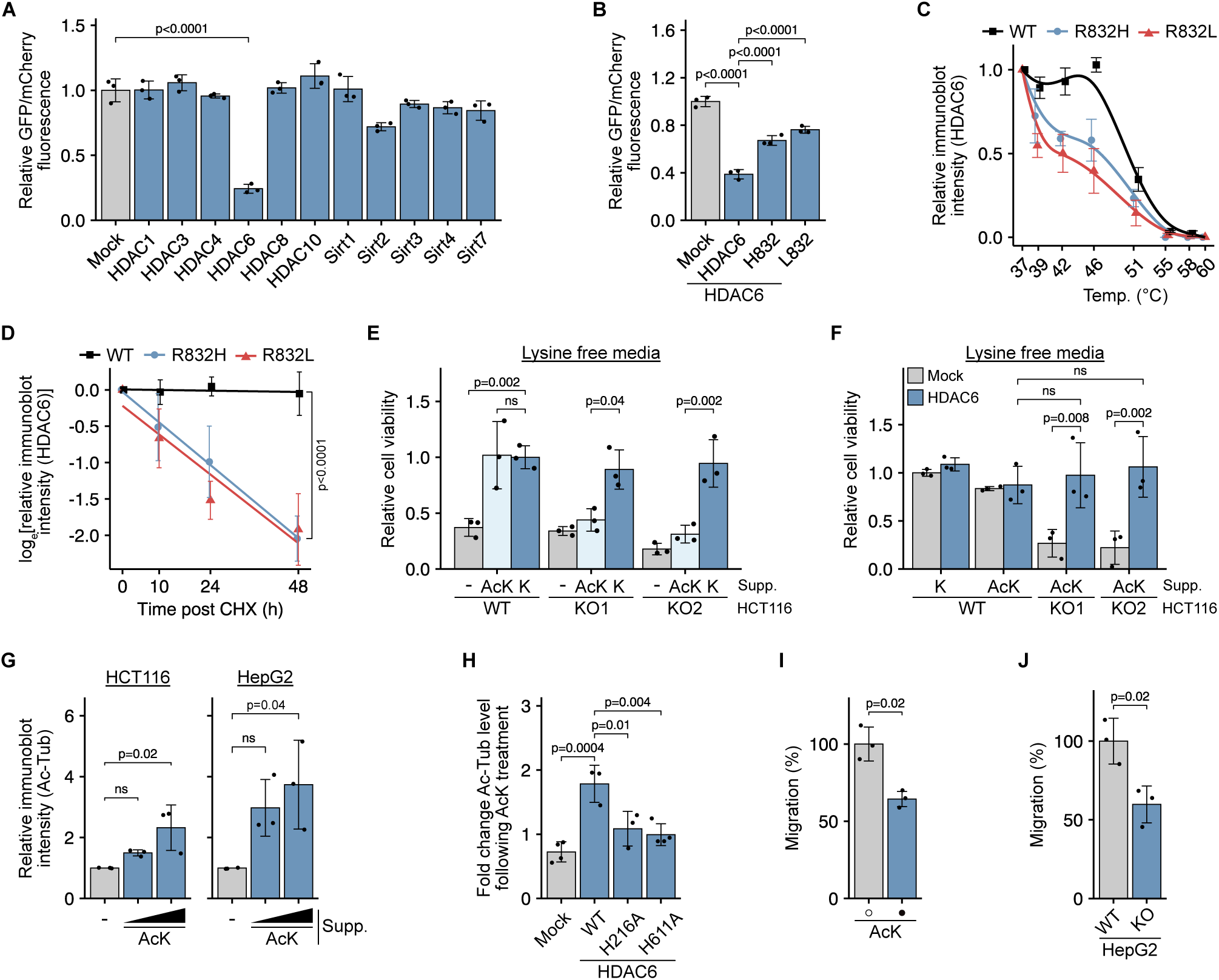
HDAC6 CD1 enables the recycling of lysine from AcK. (A) Relative GFP/mCherry fluorescence ratio measured in lysates of HEK293T cells cultured with AcK and cotransfected with mCherry-TAG-GFP encoding plasmid and indicated C-terminal Flag-tagged KDAC or empty vector. (B) Relative GFP/mCherry fluorescence ratio in lysates of HEK293T cells cultured with AcK and cotransfected with mCherry-TAG-GFP encoding plasmid and indicated HDAC6-Flag variant, or control plasmid. (C) Relative amount of WT or natural HDAC6-Flag variants quantified by anti-Flag immunoblotting, following heat denaturation at 37–61*^◦^*C. Lines serve as a guide to the eye. Data are reported as mean ± S.E.M. (D) Log relative HDAC6-Flag levels in HCT116 cells expressing WT or indicated natural HDAC6-Flag variant, described as a function of time post-CHX treatment. (E) Relative viability of WT and HDAC6-knockout HCT116 cells cultured in lysine-deficient media supplemented with 0.8 mM lysine or AcK, or without supplements. Data are presented relative to WT cells cultured with lysine. (F) Relative viability of WT and HDAC6-knockout HCT116 cells cultured in lysine-free media, as a function of lysine or AcK supplement (as described in E), and expression of HDAC6-Flag. Data are presented relative to mock-transfected WT cells cultured with lysine. (G) Relative α-tubulin K40 acetylation levels in HCT116 and HepG2 cells as a function of AcK concentration (0, 2, and 5 mM). Bars represent the ratio between AcK40 α-tubulin and α-tubulin immunoblot intensities. (H) Fold change in α-tubulin K40 acetylation levels in HDAC6-knockout HCT116 cells expressing WT or natural HDAC6-Flag variants, in response to 5 mM AcK treatment. Bars represent the fold change in AcK40 α-tubulin immunoblot intensity. (I) Effects of AcK treatment on HepG2 cell migration. Data are presented as a percentage of migrated HepG2 cells cultured without AcK. (J) Effects of HDAC6 on HepG2 cell migration. Data are presented as a percentage of migrated HepG2 cells. Data represent the mean ± S.D. of at least three independent experiments, unless indicated otherwise. Statistical analysis was performed using the Welch two-sample t-test (I, J), ANOVA followed by Tukey post hoc analysis (A, B, E, F, G, H), or pairwise trend comparisons with *p*-values adjusted for multiple comparisons using the Holm method (D). ns, non-significant.

The role of HDAC6 in AcK metabolism aligns with the association between the natural variants R832L and R832H and increased AcK levels (rs61735967; Figure S1A). However, the effect of this natural variation on AcK metabolism and the underlying mechanism are currently unknown. Residue R832 is not part of CD1 (residues 84–440) and is located between CD2 and the ubiquitin-binding domain (Figure S1B). Nevertheless, significantly higher AcK incorporation efficiency was measured in cells expressing the natural variants R832H and R832L, compared to WT HDAC6, indicating lower CD1 deacetylase activity and higher AcK cellular levels (Figure 2B). These results are in agreement with GWAS data (Figure S1A), and prior measurements showing that the activity of the isolated domains is impaired and that domains beyond the individual catalytic domain are required for full CD1 and CD2 activity.^23,39–42^

Residue R832 in HDAC6 corresponds to S795 in zHDAC6. According to the crystal structure of zCD1-CD2 (PDB: 7QNO),^43^ S795 occupies a strategic position near CD1, ‘packed’ between zCD1 and an unstructured loop linking zCD1 and zCD2 (Figure S3D). The structure further indicates a stabilizing electrostatic interaction between S795 and conserved E692 (E729 in human HDAC6). Assuming a parallel stabilizing interaction between R832 and E729, we hypothesized that mutation of R832 could destabilize HDAC6, leading to reduced levels of active HDAC6 and an apparent decrease in activity. Supporting this hypothesis, immunoblot analysis of total lysates from HEK293T cells overexpressing HDAC6 revealed significantly lower R832L HDAC6 levels, compared to WT (Figures S3E). In addition, R832L HDAC6 exhibited markedly lower residual CD1 activity after incubation at 37*^◦^*C (Figure S3F). Thermal denaturation assays further demonstrated a substantial reduction in the thermodynamic stability of both R832L and R832H mutants (Figure 2C). Consistently, in a cycloheximide (CHX) chase assay, WT HDAC6 remained stable in cells for up to 48 h, while the R832L and R832H mutants showed reduced cellular stability, with estimated half-lives of less than 20 h (Figure 2D). Overall, these findings indicate that the natural variants R832H and, more notably, R832L are less stable compared to WT HDAC6, in agreement with the increased AcK serum levels associated with this natural polymorphism.

The deacetylation of AcK by HDAC6 produces the essential amino acid lysine, providing a potential mechanism for lysine recycling. To test this, we assessed the viability of WT and HDAC6-knockout cells cultured in lysine-free media. The WT cells exhibited reduced viability in lysine-free media, but their viability improved significantly when culture media were supplemented with 0.8 mM lysine or AcK (Figures 2E and S3G). In contrast, HDAC6-knockout cells showed improved viability when media were supplemented with lysine, but not with AcK. In addition, exogenous expression of HDAC6 in HDAC6-knockout cells restored their ability to proliferate in lysine-free media supplemented with AcK (Figures 2F and S3H). Hence, the deacetylation of AcK by HDAC6 CD1 may serve as a lysine salvage mechanism to support cell metabolism and proliferation.

Given that full HDAC6 activity depends on both catalytic domains^23,41^, and that acetate is generated by both CD1 and CD2, we examined the possibility of AcK-dependent cross-domain interplay. We found that in cells, AcK supplementation resulted in significantly higher levels of α-tubulin K40 acetylation, a known HDAC6 CD2 substrate (Figures 2G, 2H, and S3I). The AcK-dependent increase in α-tubulin K40 acetylation was observed only with WT HDAC6, but not with the CD1 mutant, and was not due to differences in HDAC6 expression levels (Figure S3J), suggesting a possible mechanism involving CD1 activity. Thus, although AcK is a substrate of CD1, AcK supplementation increased α-tubulin K40 acetylation in an HDAC6-dependent manner, indicating a crosstalk between the catalytic activities of CD1 and CD2. Moreover, α-tubulin is dynamically acetylated in response to various conditions, and was suggested to regulate microtubule-dependent cell motility.^26,44–46^ Therefore, we investigated whether AcK has a role in modulating cell migration by targeting HDAC6. A transwell migration assay revealed that AcK treatment significantly reduced HepG2 cell migration (Figure 2I), consistent with the lower migration observed in HDAC6 knockout HepG2 cells (Figure 2J).

Taken together, these findings provide a mechanism for the observed association between HDAC6 polymorphism and elevated AcK levels, and show that HDAC6 CD1 can recycle AcK to produce free lysine, which is an essential amino acid. Furthermore, AcK promotes increased α-tubulin K40 acetylation and reduced cell migration.

### Functionally opposite to HDAC6 CD1, SSAT2 catalyzes lysine acetylation

In HDAC6-knockout cells, lysine and acetate supplementation increased cellular AcK levels (Figure S2L), suggesting that an additional enzyme besides HDAC6 may contribute to AcK metabolism, in agreement with our GWAS-based analysis implicating SSAT2 (Figure S1A). It was previously demonstrated, and confirmed here (Figures S4A and S4B), that SSAT2 can function as an *N* ^ε^-lysine acetyl transferase in vitro. However, little is known about the role of mammalian SSAT2 in cellular AcK synthesis. Using our GCE-based assay, we found a robust and lysine concentration-dependent increase in AcK incorporation into the GFP-N150TAG-HA and mCherry-TAG-GFP reporters upon coexpression with SSAT2 (Figures 3A, S4C, and S4D). In addition, significantly higher levels of released acetate were measured when zCD1 was incubated with metabolic extracts from HDAC6-knockout HCT116 cells overexpressing SSAT2, compared with mock-transfected cells (Figure 3B). These data show that SSAT2 can catalyze the *N* ^ε^-acetylation of free lysine in cells.

**Figure 3.**
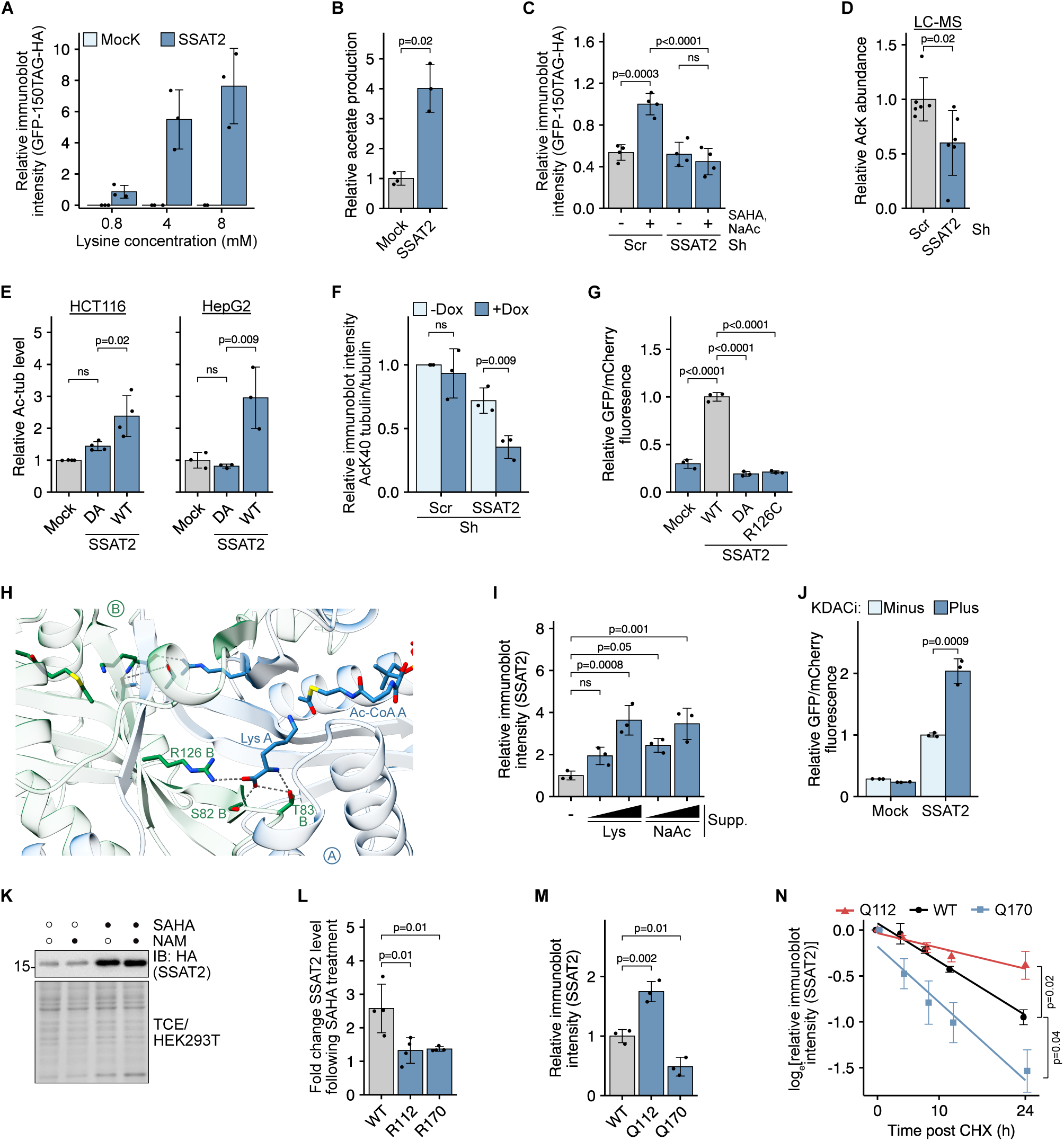
SSAT2-catalyzed AcK synthesis in cells is regulated by acetylation. (A) Relative levels of full-length GFP-HA expressed by incorporation of enodenous AcK into GFP-N150TAG-HA, in HEK293T cells cotransfected with HA-SSAT2 expressing plasmid or empty vector and cultured with increasing concentration of lysine in culture media. Bars represent relative GFP-HA immunoblot intensities. (B) Relative acetate production following incubation of recombinant zCD1 with metabolites extracted from HDAC6-knockout HCT116 cells transfected with SSAT2 expression plasmid or empty vector. (C) Relative levels of full-length GFP expressed by incorporation of enodenous AcK into GFP-N150TAG-HA, in HEK293T with stable Dox-dependent expression of Sh against SSAT2 or scrambled Sh sequence, and cultured with or without SAHA and NaAc. Bars represent relative GFP-HA immunoblot intensities. (D) Relative abundance of AcK quantified by LC-MS, in HDAC6-knockout HepG2 cells with stable Dox-dependent expression of Sh against SSAT2 or scrambled Sh sequence. (E) Relative α-tubulin K40 acetylation levels in HCT116 or HepG2 cells transfected with WT or inactive (DA) HA-SSAT2 expressing plasmid, or empty vector. Bars represent the ratio between AcK40 α-tubulin and α-tubulin immunoblot intensities. (F) Relative α-tubulin K40 acetylation levels in HepG2 cells with Dox-dependent expression of Sh against SSAT2 or scrambled sequence. Bars represent the ratio between AcK40 α-tubulin and α-tubulin immunoblot intensities as a function of Dox-induced expression of Sh. (G) Relative GFP/mCherry fluorescence ratio measured in lysates of HEK293T cells cotransfected with mCherry-TAG-GFP encoding plasmid and indicated HA-SSAT2 variant or empty vector. (H) Structural model of SSAT2 dimer with bound lysine and Ac-CoA. Monomers A and B are labeled and colored blue and green, respectively. Model was created using Boltz-2. ^38^ (I) Relative HA-SSAT2 levels in HepG2 cells expressing HA-SSAT2 and cultured without supplements or increasing amounts of lysine (4 mM and 8 mM) or NaAc (5 mM and 10 mM). Bars represent the relative HA-SSAT2 immunoblot intensities. (J) Relative GFP/mCherry fluorescence ratio measured in lysates of HEK293T cells cultured with (plus) or without (minus) KDACi (NAM and SAHA), and transfected with mCherry-TAG-GFP encoding plasmid or empty vector. (K) Immunoblotting using anti-HA antibody showing the effects of NAM and SAHA on HA-SSAT2 protein levels in HEK293T cells. (L) Fold change in levels of WT or indicated HA-SSAT2 mutant in HEK293T cells. Bars represent the fold change in HA-SSAT2 immunoblot intensities, in response to SAHA treatment. (M) Relative HA-SSAT2 levels in HEK293T cells expressing WT or indicated mutant HA-SSAT2. Bars represent the relative HA-SSAT2 immunoblot intensities. (N) Log relative HA-SSAT2 levels in HCT116 cells expressing WT or indicated mutant HA-SSAT2, described as a function of time post-CHX treatment. Data are reported as mean ± S.E.M. Data represent the mean ± S.D. of at least three independent experiments, unless indicated otherwise. Statistical analysis was performed using the Welch two-sample t-test (B, D), ANOVA followed by Tukey post hoc analysis (A, C, E, F, G, I, J, L, M), or pairwise trend comparisons with *p*-values adjusted for multiple comparisons using the Holm method (N). ns, non-significant. Scr, scrambled Sh sequence.

To determine the contribution of endogenous SSAT2 to cellular AcK levels, we created a stable doxycycline (Dox)-dependent SSAT2-knockdown HEK293T cell line, which is most suitable for GCE-based assays. UAG suppression was not affected by lysine deacetylase inhibitors (KDACi; Figure S4E), while the expression of full-length GFP-N150TAG-HA was AcK-dependent (Figure S4F), and significantly increased in scrambled Sh control cells following treatment with suberoylanilide hydroxamic acid (SAHA, an inhibitor of Zn^2+^-dependent deacetylases) and sodium acetate (NaAc), to inhibit HDAC6 and increase cellular acetyl-CoA levels, ^47,48^ respectively (Figures 3C, S4G, and S4H). In contrast, no increase in AcK incorporation was found in SSAT2-knockdown cells, suggesting that AcK synthesis is SSAT2-dependent. Finally, taking advantage of the higher levels of SSAT2 in HepG2 cells (Figure S4I), we created stable Dox-dependent SSAT2-knockdown cells on the background of the WT and HDAC6-knockout HepG2 cell lines (Figure S4J). LC-MS analysis of metabolic extracts from cells cultured in standard DMEM media found lower levels of AcK in HDAC6-knockout/SSAT2-knockdown cells, relative to scrambled control (Figure 3D). Together, these data show that although in vitro the preferred substrate of SSAT2 is thialysine, SSAT2 can contribute to elevated cellular AcK levels.

Considering the effect of AcK on α-tubulin K40 acetylation, we investigated whether SSAT2-dependent elevated cellular AcK can lead to increased α-tubulin K40 acetylation. Overexpression of WT SSAT2, but not the inactive S82D/T83A mutant (DA-SSAT2), increased α-tubulin K40 acetylation (Figures 3E and S4K) without altering endogenous HDAC6 levels (Figure S4L). Coimmunoprecipitation (co-IP) assays confirmed that SSAT2 does not stably interact with the major α-tubulin deacetylases Sirt2 and HDAC6 (Figures S4M and S4N). Consistently, we found significantly lower α-tubulin K40 acetylation in SSAT2-knockdown HepG2 cells, compared to scramble control (Figure 3F). Thus, these data show that elevated SSAT2 catalytic activity results in increased α-tubulin K40 acetylation, via a possible mechanism involving elevated AcK levels that modulate HDAC6 CD2 deacetylase activity.

Next, we inquired whether the aforementioned SNP in *SSAT2* affects SSAT2 activity. In particular, the R126C variant (rs13894-G), with a prevalence of 4.4–10.8% in the Genome Aggregation Database, is strongly associated with decreased AcK levels.^49^ In HEK293T cells cultured without AcK supplementation, we measured significantly lower incorporation of AcK into reporter proteins when coexpressed with R126C SSAT2, compared to WT SSAT2, similar to that measured for the catalytically inactive mutant DA-SSAT2 (Figures 3G and S5A). We found no difference in SSAT2 expression levels, suggesting that the R126C mutation renders SSAT2 inactive. To suggest a mechanism for the inhibitory effect of the R126C mutation, we used Boltz-2 to predict the structure of the dimeric SSAT2 in complex with lysine and Ac-CoA (Figure 3H).^38^ The suggested model predicts that the Cα amino group of the lysine substrate bound to the active site of monomer A, interacts with S82 and T83 of monomer B, explaining the lack of activity of the DA-SSAT2. Importantly, R126 of monomer B interacts with the Cα carboxylic group of the substrate bound to monomer A.

Hence, R126 plays the same role as K353 in HDAC6 CD1 in determining the enzyme’s specificity for lysine residues with a negatively charged Cα carboxylate group. Collectively, these data explain the association between the naturally occurring R126C SSAT2 variant and reduced serum AcK levels.

While measuring SSAT2 activity in cells, we found that endogenous and exogenously-expressed SSAT2 (but not GFP) are stabilized at elevated concentrations of acetate or lysine (Figures 3I, S5B, and S5C). Therefore, we hypothesized that acetylation, driven by the supplemented acetate, regulates SSAT2 levels. Supporting this, treatment with KDACi led to marked stabilization of SSAT2 (Figure S5D) and elevated cellular AcK levels, as indicated by a significantly higher GFP/mCherry fluorescence ratio measured in KDACi-treated cells expressing SSAT2 (Figure 3J). Further analyses found that KDACi treatment had no effect on the in vitro activity of SSAT2 expressed in HEK293T cells (Figure S5E), but that SSAT2 was stabilized following treatment with SAHA and not nicotinamide (NAM, an inhibitor of nicotinamide adenine dinucleotide-dependent sirtuins; Figures 3K and S5F), suggesting that acetylation-dependent stabilization of SSAT2 is regulated by Zn^2+^-dependent deacetylases, but not sirtuins.

Tandem LC-MS/MS analysis of IPed and trypsin-digested SSAT2 expressed in cells cultured with SAHA identified lysine residues at positions 10, 29, 35, 112, and 120 as acetylation sites with high confidence, and lysine 170 (the C-terminal amino acid) with medium confidence. To identify lysine residues involved in the regulation of SSAT2 stability, a Lys-to-Arg mutation was introduced in each of these positions, including K108, which is a known SSAT2 ubiquitylation site. ^50^ We found that K112R and K170R mutations largely abrogated SAHA-dependent SSAT2 stabilization (Figures 3L and S5G). Furthermore, Lys-to-Gln mutational analysis, to mimic the acetylated state, found the K112Q mutant was accumulated to significantly higher levels, while SSAT2 K170Q levels were significantly lower (Figure 3M). Supporting these observations, quantification of immunoblot intensities following CHX treatment revealed a slower degradation rate for K112Q but faster degradation for K170Q (Figure 3N). Hence, K112 and C-terminal K170 are acetylation sites on SSAT2 regulated by Zn^2+^-dependent deacetylases, and acetylation-mimic mutations suggest that K112 acetylation stabilizes SSAT2, while acetylation of K170 promotes its degradation.

### C-terminal *N* **^ε^**-lysine acetylation is a signal for SSAT2 proteosomal degradation

Following the observed degradation of SSAT2 by C-terminal K170 acetylation, we set out to investigate the role and regulation of C-terminal lysine acetylation as a degradation signal in general, and in SSAT2 in particular. Members of the GNAT family, including the closely related SSAT1, can undergo autoacetylation,^51,52^ and importantly, the GNAT family member PA4794 from *Pseudomonas aeruginosa* can acetylate C-terminal lysine residues.^53^ Therefore, we reasoned that SSAT2 may undergo autoacetylation at K170. Supporting this, we found a marked increase in acetylation of WT SSAT2, but not the K170R mutant, following in vitro incubation with acetyl-CoA, despite K170R SSAT2 being catalytically active (Figures 4A, S4A and S4B). Moreover, K170 autoacetylation was inhibited in the presence of lysine (Figure 4B), the substrate of SSAT2, possibly due to competition over binding to the active site. Thus, SSAT2 can accept a C-terminal lysine residue as a substrate, and at lower lysine concentrations, it can catalyze its autoacetylation on C-terminal K170.

**Figure 4.**
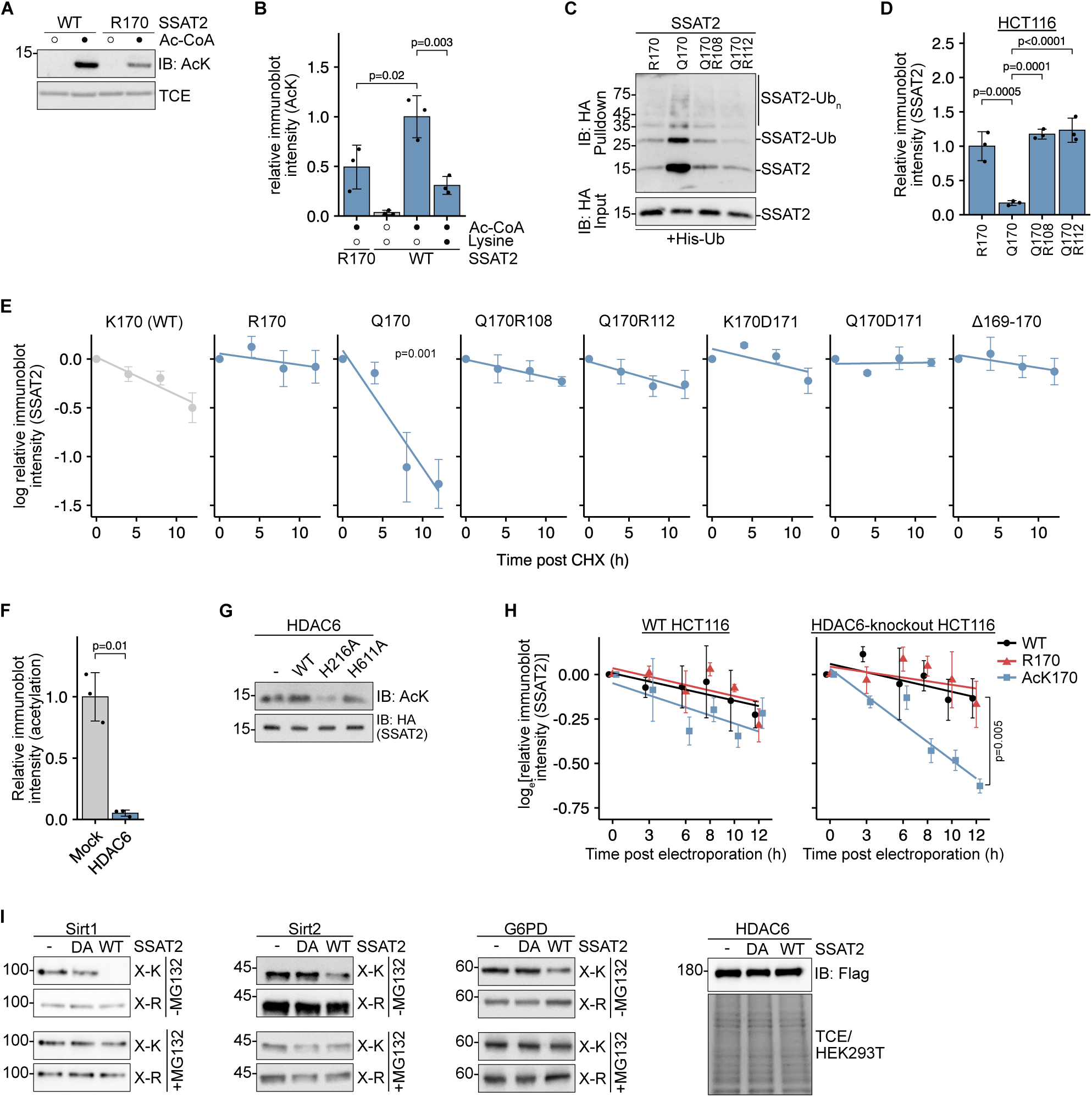
Autoacetylation of C-terminal K170 promotes SSAT2 proteosomal degradation. (A) Immunoblotting using anti-AcK antibody showing the acetylation levels of purified WT and R170 SSAT2 expressed in *E. coli*, following incubation with (•) or without (◦) Ac-CoA. (B) Acetylation levels of purified WT and R170 SSAT2 expressed in *E. coli*, following incubation with (•) or without (◦) Ac-CoA and lysine. Bars represent the relative acetylated SSAT2 immunoblot intensities. (C) Immunoblotting using anti-HA antibody showing the levels of indicated WT and mutant HA-SSAT2 in proteins isolated by cobalt resin, from lysates of HEK293T cells cotransformed with plasmid encoding HA-SSAT2 and 6×His-Ub or empty vector. (D) Levels of indicated HA-SSAT2 mutants transiently expressed in HCT116 cells. Bars represent the relative HA-SSAT2 immunoblot intensities. (E) Relative levels of WT and indicated mutant HA-SSAT2 expressed in HCT116 cells and presented as a function of time post CHX treatment. (F) Acetylation levels of AcK170 SSAT2 expressed in *E. coli*, following incubation with cleared lysates of HEK293T cells transfected with HDAC6-Flag-expressing plasmid or empty vector. Bars represent the relative K170-acetylated SSAT2 immunoblot intensities. (G) Immunoblotting using anti-AcK antibody showing the levels of AcK112 HA-SSAT2 coexpressed with WT or indicated HDAC6-Flag mutant in HEK293T cells cultured with 5 mM AcK. (H) Log normalized ratio between SSAT2 immunoblot intensity and GFP fluorescence, described as a function of time post delivery of GFP and indicated SSAT2 variant into WT and HDAC6-knockout HCT116 cells by electroporation. Data are reported as mean ± S.E.M. (I) Immunoblotting using anti-Flag antibody showing the levels of indicated C-terminal Flag-tagged proteins coexpressed with WT or inactive (DA) HA-SSAT2 in HEK293T cells with or without 5 µM MG132 treatment for 16 h. Data represent the mean ± S.D. of at least three independent experiments, unless indicated otherwise. Statistical analysis was performed using the Welch two-sample t-test (F), ANOVA followed by Tukey post hoc analysis (B, D), or pairwise trend comparisons with *p*-values adjusted for multiple comparisons using the Holm method (E, H). In-gel TCE fluorescence serves as a loading control.

To investigate the role of SSAT2 autoacetylation in regulating SSAT2 stability, we first identified the degradation pathway of K170Q SSAT2. We found that inhibition of proteosomal, but not lysosomal, degradation stabilized K170Q (Figure S6A). Furthermore, K170Q SSAT2 was more ubiquitylated compared with WT SSAT2 (Figure S6B), whereas K108R, K112R, or K170R mutants showed reduced ubiquitylation (Figure S6C). Notably, introducing K108R or K112R mutations into the K170Q mutant reduced its ubiquitylation and stabilized the protein (Figures 4C, 4D, and S6D). A CHX-chase assay further confirmed that the enhanced degradation of K170Q SSAT2 involves K108/K112 and, critically, revealed that degradation is promoted only when the acetylation-mimic mutation occurs at the ultimate C-terminal residue (Figure 4E); Q170 SSAT2 was significantly less stable relative to WT or K170Q SSAT2 that were extended by a C-terminal Asp residue (K170-D171 or Q170-D171, respectively), or truncated SSAT2 missing the C-terminal Gly-Lys residues (Δ169-170). Together, these data suggest that autoacetylation of C-terminal K170 acts as a degradation signal, promoting SSAT2 degradation via K108/K112 ubiquitylation, whereas acetylation at K112 stabilizes SSAT2 by preventing this ubiquitylation.

Given the stabilizing effect of SAHA and HDAC6’s capacity to deacetylate lysine residues at the C-terminus (via CD1) and internally (via CD2), we next asked whether AcK112 and AcK170 SSAT2 are HDAC6 substrates. We found that in vitro, both zCD1 and full-length HDAC6 catalyze the deacetylation of recombinant AcK170 SSAT2 (Figures S6E and 4F). In addition, AcK112 was deacetylated in vitro by WT HDAC6 (Figure S6F), and in cells by CD1-inactive HDAC6 (Figure 4G). These findings suggest that HDAC6 exerts opposing effects on SSAT2 stability: CD1-dependent deacetylation of C-terminal AcK170 may suppress SSAT2 degradation, whereas CD2-dependent deacetylation of K112 can promote SSAT2 degradation by enabling K108/K112 ubiquitylation. Given that the impact of K170 acetylation on SSAT2 degradation depends on K112 ubiquitylation, the deacetylation of AcK112, regulated by CD2, appears to be the critical determinant of SSAT2 turnover. Supporting this, CHX-chase assays showed that wild-type SSAT2 degraded faster in WT HCT116 cells than in HDAC6-knockout cells (Figure S6G), emphasizing the net destabilizing effect of HDAC6. Consistently, HDAC6 knockout increased levels of WT SSAT2 but not K112Q SSAT2 (Figure S6H). Hence, AcK170 and AcK112 are substrates of HDAC6 CD1 and CD2, respectively, and HDAC6 enables the proteasomal degradation of SSAT2 primarily through CD2-dependent deacetylation of AcK112.

The hypothesis that C-terminal lysine acetylation functions as a degradation signal, inferred from a Lys-to-Gln substitution, raises two key questions: first, whether glutamine is a valid mimic of AcK, given that C-terminal glutamine is a known C-degron;^54,55^ and second, whether C-terminal lysine acetylation can drive degradation of proteins beyond SSAT2.

To determine whether C-terminal acetylation, rather than the presence of a terminal glutamine, serves as a degradation signal for SSAT2, we used electroporation to simultaneously deliver purified GFP (as a normalization control) and recombinant WT, R170, or AcK170 SSAT2 into WT and HDAC6-knockout HCT116 cells. SSAT2 clearance relative to GFP fluorescence showed no significant difference between WT and R170 SSAT2 in either cell line (Figures 4H). Consistent with results observed for the K170Q mutant, AcK170 SSAT2 degraded rapidly in HDAC6-knockout cells. However, in WT cells, where AcK170 could be efficiently deacetylated, AcK170 SSAT2 was as stable as WT SSAT2 (Figure S6I). These findings provide direct evidence that C-terminal acetylation promotes SSAT2 degradation while being a substrate of HDAC6. Moreover, although C-terminal glutamine has been implicated as a C-degron targeted by Trim7,^54,55^ coexpression of WT, R170, or Q170 SSAT2 with either WT Trim7 or the inactive mutant S499A had no impact on SSAT2 levels (Figure S6J). Thus, degradation of Q170 SSAT2 proceeds via a Trim7-independent mechanism, distinct from that previously described for proteins with a C-terminal glutamine. Combined, these results establish C-terminal acetylation as a bona fide degradation signal for SSAT2, regulated by HDAC6.

To assess whether C-terminal acetylation functions as a degradation signal beyond SSAT2, we examined C-terminal Flag-tagged proteins, leveraging the fact that the canonical Flag tag terminates with a lysine (DYKDDDDK). Coexpression with WT SSAT2 led to a marked reduction in the levels of C-terminal Flag-tagged Sirt1, Sirt2, and G6PD (labeled X-K) relative to either catalytically inactive DA-SSAT2 or empty vector controls (Figure 4I). This effect was largely abolished by substituting the terminal lysine with arginine (labeled X-R), indicating that the C-terminal lysine is required for SSAT2-dependent degradation. In contrast, levels of C-terminal Flag-tagged HDAC6 remained unaffected, likely due to its deacetylase activity that can counteract SSAT2-mediated acetylation. Moreover, inhibition of proteosomal degradation using MG132 restored levels of the Flag-tagged proteins (Figure 4I) and led to the accumulation of ubiquitylated G6PD-Flag in the presence of wild-type SSAT2 (Figure S6K). These findings indicate that SSAT2 may promote proteasomal degradation of diverse proteins bearing a C-terminal lysine, thereby linking the cell’s metabolic status (AcK levels) to the proteasomal degradation pathway.

Together, the presented data show that SSAT2 C-terminal autoacetylation functions as a substrate-dependent autoregulatory degron that promotes SSAT2 proteasomal degradation, and suggest that C-terminal acetylation may serve as a degradation signal for other proteins.

## DISCUSSION

Our biochemical, cellular, and in vivo analyses reveal a coordinated system balancing lysine and AcK homeostasis through the opposing enzymatic activities of HDAC6 CD1 and SSAT2. Based on these observations, we propose a metabolic feedback mechanism that integrates protein acetylation and degradation with lysine homeostasis (Figure 5). The equilibrium between lysine and AcK is influenced by the availability of acetate and Ac-CoA; a central metabolite that also acts as a signaling molecule via lysine acetylation.^56,57^ SSAT2 autoacetylation of C-terminal K170 acts as a degradation signal, but is inhibited by high concentrations of lysine, creating a substrate-driven stabilization mechanism. The other SSAT2 substrate, Ac-CoA, can contribute to both SSAT2 destabilization (K170 acetylation) and SSAT2 stabilization (K112 acetylation). HDAC6 acts as the master regulator of this axis: its CD1 activity not only lowers free AcK levels through deacetylation but also promotes AcK synthesis indirectly, by stabilizing SSAT2; however, CD2-mediated deacetylation of AcK112 promotes SSAT2 degradation by enabling K108/K112 ubiquitylation. This intricate architecture, characterized by feedback loops and dual-functional nodes, is an example of a regulatory mechanism designed for maintaining proteostatic and metabolic stability.

**Figure 5.**
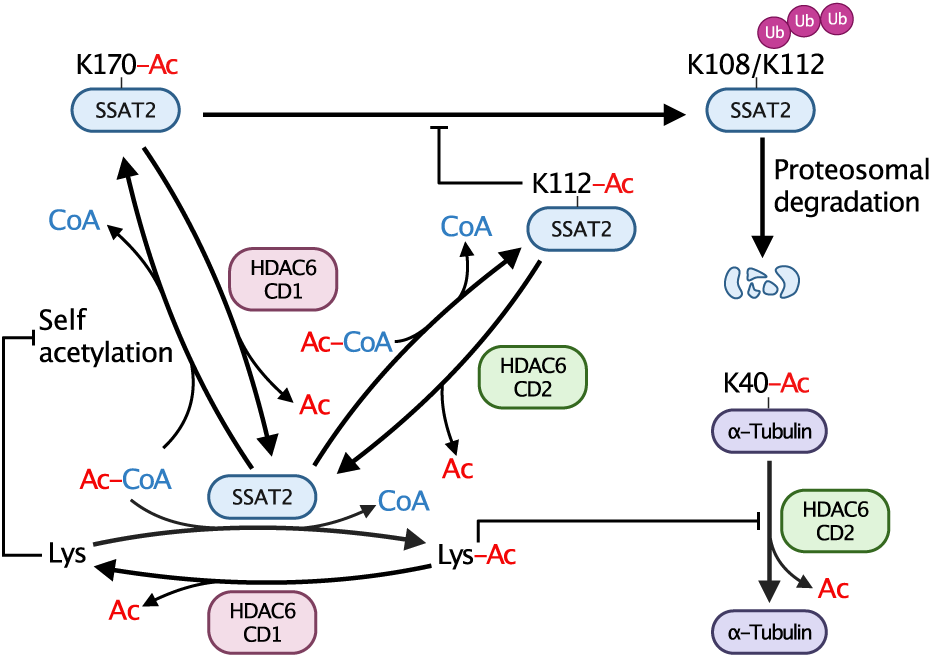
Schematic model summarizing the findings described in this manuscript regarding the regulation of lysine and AcK homeostasis by HDAC6 and SSAT2.

Similar to SSAT2 and HDAC6 CD1, SSAT1, the homolog of SSAT2, and HDAC10, the other class IIb HDAC, have opposing catalytic activities; the acetylation and deacetylation of spermidine, respectively^58–60^. We revealed a mechanism for the mutual regulation of SSAT2 and HDAC6 activity, but a reciprocal mechanism for SSAT1 and HDAC10 has not been identified. In addition, our findings highlight a relatively understudied topic: the metabolic fate of free post-translationally modified amino acids. For instance, phosphoserine is a well-known metabolite,^61^ and trimethylated lysine can be metabolized to L-carnitine in multiple steps.^62^ However, it remains to be determined which amino acids conjugated to a PTM exist in free form within the cellular environment, whether they have functional roles, and how they are metabolized.

While AcK has long been documented as a metabolite, its physiological significance remained poorly understood. Our data demonstrate that the equilibrium between AcK and lysine, mediated by HDAC6 CD1 and SSAT2, may function as a lysine-conserving mechanism. Lysine catabolism is specific and prominently involves *N* ^ε^-conjugation.^63^ Thus, *N* ^ε^-acetylation can mask the ε-amino group and form a pool of protected lysine that can be converted back to lysine. Moreover, animals fed a lysine-restricted diet lose weight more slowly than those fed a diet restricted in other essential amino acids,^64,65^ suggesting the existence of a specialized metabolic buffer such as AcK and the HDAC6/SSAT2 axis. Interestingly, the ability to synthesize lysine was lost in the metazoan lineage, and HDAC6, with its two tandem HDAC domains, may have conferred a meaningful adaptive advantage.

Beyond its role in lysine conservation, we found that AcK is a negative modulator of CD2, leading to increased α-tubulin acetylation and attenuated cell migration. Previous studies found that high AcK levels are associated with aging and poor sleep quality.^7,8^ In addition, low HDAC6 activity (by knockout or inhibition)^66–68^ or higher AcK levels^12,69^ are associated with reduced symptoms of depression. By defining AcK as a substrate of HDAC6 CD1 and a negative modulator of HDAC6 CD2, we can now suggest an overlooked link between studies focusing on HDAC6 activity and those measuring AcK levels.

Our findings provide an additional perspective for interpreting prior studies. It was recently suggested that AcK can be cotranslationally incorporated into proteins by the lysyl tRNA synthetase/tRNA pair.^15^ Some conclusions relied on experiments based on AcK supplementation, including isotopically labeled AcK, where AcK deacetylation by HDAC6 CD1 should be considered. In another study, SSAT2 was reported to promote the degradation of the α subunit of hypoxia-inducible factor 1 (HIF-1α) and suppress HIF-1 transcriptional activity, by stabilizing protein-protein interactions.^36^ Notably, these effects were partly dependent on SSAT2’s catalytic activity, potentially due to AcK production that inhibits HDAC6, whose deacetylase activity stabilizes and activates HIF-1α under hypoxia.^70^

Another key finding of this study is the identification of C-terminal lysine acetylation as a degradation signal of SSAT2, and potentially other proteins. The C-terminal autoacetylation of SSAT2 is inhibited by lysine, providing a substrate-driven stabilizing mechanism, and warrants a follow-up study to evaluate SSAT2 activity in patients with hyperlysinemia. In addition, our observation that C-terminal lysine acetylation promotes the degradation of proteins beyond SSAT2 suggests a broader regulatory mechanism. However, it remains to be determined whether C-terminal lysine acetylation is a general C-terminal degradation signal (C-degron). If so, the SSAT2/HDAC6 CD1 pair can theoretically function as an ‘On-Off’ switch for any protein harboring a C-terminal lysine, allowing the cell to couple protein half-life directly to its metabolic energetic state.

Collectively, these insights outline a unifying model in which HDAC6 and SSAT2 orchestrate reversible acetylation events that couple metabolite recycling with protein quality control, thereby linking metabolic homeostasis to protein post-translational regulation.

## Experimental Procedures

### Materials and Reagents

The full inventory of commercially available compounds purchased from Thermo Fisher, Sigma, and Biological Industries is described in the Materials table.

### Generation of plasmid constructs

To express Flag-tagged HDAC6 and HA-tagged SSAT2 mutants in mammalian cells, site-directed mutagenesis was performed using pCDNA3.1/HDAC6-Flag (Addgene plasmid #13823) and pCDNA3.1/HA-SSAT2 (Genscript) as templates, respectively. The resulting constructs included HDAC6 variants (H216A, H611A, H216A/H611A, R832H, and R832L) and SSAT2 variants (R10, R29, R35, R108, R112, R120, R170, Q170, Q170/R108, Q170/R112, Q112, R126C, and S82D/T83A). Truncated versions of *HDAC6* were generated via inverse PCR followed by self-ligation. For mammalian expression of HDAC10, the *HDAC10* cDNA was PCR-amplified from pACYC/HDAC10-Flag^71^ and subcloned into the pcDNA3.1 backbone between the AflII and XbaI restriction sites. For the site-specific, cotranslational incorporation of AcK into amber (TAG) mutants of HA-SSAT2, mCherry-GFP-HA, and GFP-HA, an orthogonal translation system was employed in cultured mammalian cells.^72,73^ This system utilized a previously described pBudCE4.1-based single vector carrying an orthogonal evolved pyrrolysyl-tRNA synthetase/amber suppressor tRNA pair.^74^ In-frame TAG mutants of *SSAT2* were generated by overlapping PCR and subsequently ligated into the pBud vector between NotI and XhoI sites, downstream of the EF1α promoter. To facilitate the incorporation of alternative non-canonical amino acids, analogous vectors were generated by cloning the respective evolved synthetases between BamHI and EcoRI restriction sites.

For recombinant expression of HDAC6 in *E. coli*, the gene fragment encoding residues 84–835 was amplified from pcDNA3.1/HDAC6-Flag (mentioned above) and subcloned into the pSH21 backbone between NheI and NotI restriction sites. The resulting construct expressed HDAC6 fused to an N-terminal 6×His-tagged maltose-binding protein (MBP) followed by a tobacco etch virus (TEV) protease cleavage site. Catalytic mutants (H216A and H611A) were generated using site-directed mutagenesis. The pET28(+)/zCD1 plasmid, encoding the first catalytic domain of zCD1 (residues 60–419), was a gift from David Christianson^23^; the K330E zCD1 mutant was subsequently generated via site-directed mutagenesis. To produce tag-free WT and site-specifically acetylated SSAT2, an intein-mediated protein splicing strategy was employed. ^75^ The pETDuet/SSAT2-TKPl plasmid (Genscript), encoding WT SSAT2 fused to a C-terminal temperature-sensitive intein, MBP, and a 6×His tag, served as the primary template. Lys-to-Arg mutations and in-frame TAG mutations were introduced by site-directed mutagenesis. For genetic code expansion, these *SSAT2* variants were PCR-amplified and cloned into a pCDF vector between NcoI and NotI restriction sites. This vector carries the gene for the U25C mutant of PylT under the control of the lpp promoter.

All plasmid constructs were verified by Sanger sequencing. Detailed information regarding plasmids, including Addgene accession numbers and source identifiers, is provided in the Materials Table.

### Cell culture

HEK293T, HepG2, and HCT116 cell lines were maintained in a humidified incubator at 37*^◦^*C with 5% CO_2_. HEK293T and HepG2 cells were cultured in Dulbecco’s Modified Eagle Medium (DMEM) supplemented with 10% (v/v) heat-inactivated fetal bovine serum (FBS), 2 mM L-glutamine, 1 mM sodium pyruvate, 100 U/mL penicillin G sodium, and 100 µg/mL streptomycin sulfate. HCT116 cells were maintained in McCoy’s 5A medium supplemented with 10% (v/v) heat-inactivated FBS, 2 mM L-glutamine, 1 mM sodium pyruvate, 100 U/mL penicillin G sodium, and 100 µg/mL streptomycin sulfate. Where indicated, endogenous KDACs were inhibited by the addition of 15 mM NAM and 3 µM SAHA to culture media. All cell lines were routinely screened for mycoplasma contamination using PCR-based assays.

### Generation of CRISPR/Cas9 knockout cell lines

To generate *HDAC6* knockout models, HCT116 and HepG2 cell lines were engineered using the CRISPR/Cas9 system. Cells were transfected with pX459 vectors encoding sgRNAs targeting human *HDAC6* (pX459/sgRNA1 or pX459/sgRNA2; gifts from Jun Zhang)^76^ using Lipofectamine 2000 according to the manufacturer’s instructions. Following transfection, cells were subjected to antibiotic selection to enrich for Cas9-expressing populations. HCT116 cells were treated with 1 µg/mL puromycin, and HepG2 cells were treated with 1.8 µg/mL puromycin for 72–96 h. After the selection period, cells were allowed to recover in puromycin-free growth medium for 48 h. Single-cell clones were subsequently isolated via limiting dilution in 96-well plates. After 14–21 days of expansion, individual colonies were trypsinized and transferred to larger culture formats. Successful knockout of *HDAC6* was validated in expanded clones by immunoblotting for the loss of HDAC6 protein expression. Confirmed knockout clones were further expanded and maintained as described in the cell culture section.

### Stable SSAT2-knockdown cell lines

To generate stable knockdown models, HEK293T and HepG2 cells were transduced with lentiviral vectors expressing either an shRNA targeting human *SSAT2* (5’-GGGCTATGGGATATACTATTT-3’) or a non-targeting scrambled control sequence (5’-CAACAAGATGAAGAGCACCAA-3’). Lentiviral particles were produced in HEK293T cells by co-transfection of the lentiviral transfer vector together with four helper plasmids encoding Gag/Pol, Rev, Tat, and VSV-G using polyethylenimine (PEI). Transductions were performed in growth medium supplemented with 1 µg/mL polybrene. At 24 h post-transduction, the viral supernatant was replaced with fresh culture medium. Selection was initiated 48 h post-transduction by the addition of 2 µg/mL puromycin. Selection was maintained for 72 h, and the resulting stable cell populations were expanded and maintained under puromycin selection. For experiments requiring gene silencing, *SSAT2* knockdown was triggered by the addition of 5 µg/mL Dox to the culture medium for 72 h.

### In cellulo HDAC6 CD1 activity assay

HEK293T or HCT116 cells were seeded in 24-well plates at a density of 1×10^5^ cells per well and cultured for 24 h. Prior to transfection, the culture medium was replaced with fresh complete DMEM or McCoy’s 5A supplemented with 2 mM AcK. Following 30 min of equilibration, cells were co-transfected with 0.75 µg of plasmid DNA encoding HDAC6 variants and either GFP-N150TAG-HA or mCherry-TAG-GFP-HA reporter constructs. Transfection of HEK293T cells was performed using PEI at a 3:1 (w/w) PEI:DNA ratio, while HCT116 cells were transfected using Lipofectamine 2000 according to the manufacturer’s protocol. For small-molecule inhibition assays, compounds were added to the HEK293T culture media at the indicated concentrations 5 h post-transfection, alongside 1 mM AcK. Cells were incubated for an additional 48 h prior to harvesting. Cells were washed twice with ice-cold PBS and lysed in RIPA buffer [50 mM Tris-HCl, pH 8.0, 150 mM NaCl, 1% (v/v) Triton X-100, 0.5% (w/v) sodium deoxycholate, and 0.1% (w/v) SDS] supplemented with DNAse I and protease inhibitor cocktail (1.2 µg/mL leupeptin, 1 µM pepstatin A, 100 µM phenylmethanesulfonyl fluoride (PMSF), and 1 µg/mL aprotinin). Lysates were clarified by centrifugation at 15,000×g for 10 min at 4*^◦^*C, and total protein concentration was determined using the BCA Protein Assay Kit. To measure reporter fluorescence, 10 µg (HEK293T) or 30 µg (HCT116) of total protein lysate were transferred to a 96-well black-walled plate and adjusted to a final volume of 100 µL with PBS. Fluorescence intensity was quantified using a Spark microplate reader (Tecan, Männedorf, Switzerland) with the following settings: excitation/emission at 485/535 nm for GFP and 580/620 nm for mCherry.

### Cell viability assay

Cell viability and metabolic activity were quantified using the Cell Counting Kit-8 (CCK-8), utilizing 2-(2-methoxy-4-nitrophenyl)-3-(4-nitrophenyl)-5-(2,4-disulfophenyl)-2H-tetrazolium monosodium salt (WST-8), according to the manufacturer’s protocol. To assess the impact of AcK on the proliferation of WT and HDAC6-knockout cell lines, cells were seeded in 96-well plates at a density of 1×10^4^ cells per well in complete DMEM. After 24 h, the growth medium was removed, and cells were washed three times with PBS. The medium was then replaced with lysine-free complete medium (#280001200; Silantes, Mering), formulated to match the standard growth medium composition excluding L-lysine. The lysine-free medium was supplemented with either 0.8 mM L-lysine (positive control) or 0.8 mM AcK (AcK medium), as required. Following incubation for the indicated time points, 10 µL of CCK-8 reagent was added to each well containing 100 µL of medium. After a 2 h incubation at 37*^◦^*C, the absorbance was measured at 450 nm, with a reference background subtraction at 600 nm, using a Spark microplate reader.

### CHX chase assay

To evaluate the half-life of HDAC6 and SSAT2 variants, HCT116 cells were seeded in 24-well plates at a density of 1×10^5^ cells per well. After 24 h, cells were transiently transfected with the indicated plasmids using Lipofectamine 2000. At 5 h post-transfection, the medium was replaced with fresh complete McCoy’s 5A. Following an additional 24 h incubation, de novo protein synthesis was inhibited by the addition of 50 µg/mL CHX. Cells were harvested at specified time intervals (0–48 h) post-treatment. Collected cells were washed with ice-cold PBS and lysed in PBS supplemented with 1% (v/v) Triton X-100, DNAse I, and protease inhibitor cocktail. Lysates were clarified by centrifugation at 15,000×g for 10 min at 4*^◦^*C. Total protein concentration was determined by BCA assay, and 30 µg of total protein per sample was resolved by SDS-PAGE. The steady-state levels of HDAC6 or SSAT2 were analyzed by Western blotting, and degradation kinetics were determined by normalizing target protein levels to endogenous enolase.

### Cellular thermal shift assay

HEK293T cells transiently expressing HDAC6 were washed once with PBS and resuspended in PBS at 6×10^6^ cells/mL. 50 µL of the suspension was aliquoted into PCR tubes and heated in a gradient PCR thermal cycler at the indicated temperatures (37–60*^◦^*C) for 3 min. Samples were immediately equilibrated at 25*^◦^*C for 3 min. For lysis, 40 µL of cell suspension was mixed 1:1 with 2× RIPA buffer, incubated on ice for 20 min, and clarified by centrifugation at 15,000×g for 10 min at 4*^◦^*C to remove debris and insoluble aggregates. 60 µL of the cleared lysate was used for subsequent Western blot analysis.

### In-cell ubiquitination assay

In-cell ubiquitination assays were performed as described previously^37^ with minor modifications. Briefly, HEK293T cells were transiently transfected with the indicated expression constructs together with a His-tagged ubiquitin (His-Ub) plasmid. At 36 h post-transfection, cells were treated with 5 µM MG132 overnight, washed with ice-cold PBS, and lysed in RIPA buffer supplemented with protease inhibitor cocktail and DNase I. Lysates were clarified by centrifugation at 4*^◦^*C, and clarified lysates were incubated with cobalt resin at 4*^◦^*C overnight with rotation. Beads were washed three times with wash buffer (50 mM sodium phosphate, pH 7.4, 300 mM NaCl, 10 mM imidazole), and His-ubiquitinated proteins were eluted with elution buffer (50 mM sodium phosphate, pH 7.4, 300 mM NaCl, 150 mM imidazole) and analyzed by immunoblotting as indicated. For Flag-based immunoprecipitation, clarified lysates were incubated with anti-Flag M2 magnetic beads at 4*^◦^*C with rotation. Beads were washed with PBS, and bound proteins were eluted in 1× non-reducing Laemmli sample buffer at 70*^◦^*C for 10 min. Eluates were resolved by SDS-PAGE and subjected to immunoblot analysis using the indicated antibodies.

### Transwell migration assay

Cell migration was assessed using Transwell inserts (24-well format, 8 µm pore size). HepG2 cells were harvested, resuspended in serum-free DMEM, and seeded into the upper chambers at a density of 1.5×10^4^ cells per well in 300 µL. The lower chambers were filled with 600 µL complete DMEM containing 10% FBS as a chemoattractant. For AcK treatment, 5 mM AcK was added to both the upper and lower chambers to maintain equivalent concentrations throughout the assay. Cells were incubated for 72 h, and non-migrated cells on the upper surface of the membrane were gently removed with a cotton swab. Cells that had migrated to the lower surface were fixed with 25% methanol for 10 min at room temperature and stained with 1% (w/v) crystal violet for 10 min. Membranes were washed extensively with distilled water to remove excess dye. For quantitative analysis, stained cells were solubilized in 500 µL 10% (v/v) acetic acid, and absorbance at 595 nm was measured using a Spark microplate reader.

### Bacterial expression and purification of recombinant proteins

#### Purification of recombinant HDAC6

Recombinant human HDAC6 CD1-CD2 (aa84–835; CD12), H216A CD12, and H611A CD12 were expressed and purified as previously described^23^ with minor modifications. All constructs were transformed into *E. coli* BL21(DE3) cells for protein production. An overnight starter culture was diluted 1:100 into 6 L 2×TY medium supplemented with 100 µg/mL ampicillin and grown at 37*^◦^*C with shaking until OD_600_ reached 0.5. ZnSO_4_ was added to a final concentration of 200 µM, and the incubation temperature was reduced to 16*^◦^*C. After 30 min equilibration, protein expression was induced with 75 µM isopropyl β-L-1-thiogalactopyranoside (IPTG), and cultures were incubated for an additional 18 h at 16*^◦^*C. Cells were harvested by centrifugation at 4,000×g for 15 min at 4*^◦^*C, washed with ice-cold PBS, and stored at-80*^◦^*C. Cell pellets were resuspended in Ni-NTA binding buffer [50 mM sodium phosphate (pH 8.0), 300 mM NaCl, 5 mM β-mercaptoethanol (β-ME), 10 mM imidazole, 10% glycerol] supplemented with 0.5 mg/mL lysozyme, 20 µg/mL DNase I, and protease inhibitor cocktail. Cells were lysed by sonication on ice, and lysates were clarified by centrifugation at 10,000×g for 60 min at 4*^◦^*C. The supernatant was filtered through a 0.45 µm membrane prior to chromatography. Clarified lysates were applied to a 5 mL HisTrap HP column (Cytiva) equilibrated in binding buffer. Bound proteins were eluted with Ni-NTA elution buffer [50 mM sodium phosphate (pH 8.0), 300 mM NaCl, 300 mM imidazole, 5 mM β-ME, 5% glycerol]. Peak fractions were diluted two-fold with MBP binding buffer [20 mM Tris-HCl (pH 8.0), 100 mM NaCl, 10 mM β-ME, 5% glycerol] and loaded onto a 5 mL MBPTrap HP column (Cytiva). The His-MBP affinity tag was removed by on-column cleavage with tobacco etch virus (TEV) protease. Eluted, tag-free HDAC6 was further purified by size-exclusion chromatography using a HiLoad 26/600 Superdex 200 column (Cytiva) pre-equilibrated in SEC buffer [50 mM HEPES (pH 7.5), 100 mM KCl, 5 mM dithiothreitol (DTT), 5% glycerol]. Fractions corresponding to monomeric HDAC6 were pooled and concentrated to 1.1-3 mg/mL using a centrifugal concentrator with a 30 kDa molecular weight cutoff. Proteins were aliquoted, flash-frozen in liquid nitrogen, and stored at-80*^◦^*C. Protein purity was assessed by SDS-PAGE followed by trichloroethanol (TCE)-based in-gel fluorescence visualization (as detailed below for the general Western blotting analysis procedure). Protein identity was confirmed by immunoblotting using HDAC6-specific antibodies.

#### Purification of recombinant zCD1

Wild-type and K330E zCD1 were expressed and purified as described above for HDAC6 CD12, with the following modifications. Constructs were transformed into *E. coli* BL21(DE3), and bacterial cultures were grown in 2×TY medium supplemented with kanamycin. Protein expression was induced, and cells were harvested as described for HDAC6. Following Ni-affinity purification and on-column TEV protease cleavage to remove the His-MBP tag, the eluted, untagged zCD1 protein was subjected to an additional ion-exchange chromatography (IEC) step prior to size-exclusion chromatography. Specifically, the TEV-cleaved protein solution was diluted 1:10 in ice-cold IEC buffer [20 mM Tris-HCl (pH 8.0), 10 mM β-ME, 5% glycerol] to reduce ionic strength and loaded onto a 5 mL HiTrap Q FF column (Cytiva) pre-equilibrated in IEC buffer. Bound protein was eluted using a linear gradient to IEC elution buffer [20 mM Tris-HCl (pH 8.0), 10 mM β-ME, 5% glycerol, 1 M NaCl] over 20 column volumes (CV). Fractions containing zCD1, as determined by SDS–PAGE analysis, were pooled and further purified by size-exclusion chromatography on a Superdex 200 column equilibrated in SEC buffer [50 mM HEPES (pH 7.5), 100 mM KCl, 5 mM DTT, 5% glycerol].

#### Purification of recombinant WT and acetylated SSAT2

The incorporation of AcK into the C-terminal position is expected to yield a truncated protein lacking the last amino acid, which is essentially indistinguishable from the full-length protein. To resolve this, SSAT2 was engineered as a fusion protein containing a C-terminal temperature-sensitive intein followed by MBP and a 6×His tag. This strategy allowed purification of the full-length precursor protein in the first stage and subsequent release of native SSAT2 as the N-extein through controlled intein-mediated cleavage.

For the expression of AcK170 SSAT2, *E. coli* BL21(DE3) cells were co-transformed with the pBK plasmid encoding the evolved AcKRS3 synthetase and the pCDF/TAG170 SSAT2-TKPl plasmid harboring the *SSAT2* gene with a TAG mutation at codon 170. Overnight cultures grown in LB containing 50 µg/mL spectinomycin and 50 µg/mL kanamycin were diluted to OD_600_=0.05 in 3 L 2×TY supplemented with the same antibiotics and incubated at 37*^◦^*C with shaking (130 rpm). At OD_600_=0.4, the medium was supplemented with 10 mM AcK and 20 mM NAM. When OD_600_ reached 1.2, ZnSO_4_ was added to 200 µM, the temperature was reduced to 16*^◦^*C, and after 1 h equilibration, protein expression was induced with 0.5 mM IPTG. Cultures were grown for an additional 24 h at 16*^◦^*C. Cells were harvested and lysed in ice-cold lysis buffer [50 mM sodium phosphate (pH 8.0), 300 mM NaCl, 10 mM imidazole]. Lysates were clarified by centrifugation at 10,000 × g for 60 min at 4*^◦^*C, and the supernatant was applied to tandem 5 mL HisTrap FF columns pre-equilibrated in lysis buffer in the cold room. After washing with 20 CV of ice-cold lysis buffer, columns were equilibrated with 5 CV intein cleavage buffer [50 mM Tris-HCl (pH 8.0), 100 mM NaCl, 5% glycerol]. On-column intein cleavage was induced by incubation at 37*^◦^*C for 18 h. The cleaved, tag-free AcK170 SSAT2 was eluted with intein cleavage buffer and further purified by size-exclusion chromatography in SEC buffer [20 mM Tris-HCl (pH 8.0), 10% glycerol]. Peak fractions were pooled, concentrated as needed, flash-frozen, and stored at-80*^◦^*C.

Wild-type and R170 SSAT2 were expressed and purified using the same intein-based strategy with minor modifications. BL21(DE3) cells were transformed with pETDuet/SSAT2-TKPl or pETDuet/R170 SSAT2-TKPl and cultured in 2×TY medium supplemented with 100 µg/mL ampicillin. Expression, lysis, affinity purification, intein cleavage, and size-exclusion chromatography were performed as described above.

### In vitro biochemical assays

#### In vitro deacetylase activity of zCD1

In vitro deacetylation assays were performed using purified recombinant WT or E330 zCD1 as described below. Reactions were carried out with a final enzyme concentration of 0.25 µM, and substrates were added to a final concentration of 6 mM and included D-isomer of *N* ^ε^-acetyllysine, L-isomer of *N* ^ε^-acetyllysine (AcK), or *N* ^α^-acetyl-L-lysine, as indicated. Reactions were incubated at 37*^◦^*C for 1 h, and at the indicated time point, 60 µL of reaction mixture was collected and quenched by addition of 2.5 µL 10% (v/v) HCl to terminate enzymatic activity, neutralized with 15 µL freshly prepared 8% (w/v) sodium bicarbonate, and clarified by centrifugation at 12,000×g for 5 min at room temperature. Acetate production was quantified using a commercially available acetic acid assay kit (NZYtech, cat#AK00081) according to the manufacturer’s protocol. In this coupled enzymatic assay, acetate concentration is proportional to the amount of NADH generated through sequential enzymatic reactions. NADH formation was monitored by measuring absorbance at 340 nm at 2 min intervals until the signal plateaued.

Steady-state kinetic parameters were determined using a discontinuous, time-course–based deacetylation assay. Reactions were performed in AcK deacetylation assay buffer [20 mM HEPES (pH 7.5), 100 mM NaCl, 5 mM KCl, 1 mM MgCl_2_] in a final volume of 600 µL. Recombinant zCD1 was used at a final concentration of 0.25 µM and incubated with increasing concentrations of AcK (0.25–12 mM) at 37*^◦^*C. To ensure measurements within the linear range of product formation, 60 µL aliquots were withdrawn at 2–3 min intervals over a 0–30 min time course and quenched immediately with 2.5 µL 10% (v/v) HCl. Samples were maintained on ice until completion of the time course. Reactions were subsequently neutralized with 15 µL freshly prepared 8% (w/v) sodium bicarbonate and clarified by centrifugation at 12,000×g for 5 min at room temperature. For acetate quantification, 35 µL of the cleared supernatant was transferred to a clear, flat-bottom 96-well plate and analyzed using an acetate detection kit (NZYtech) according to the manufacturer’s instructions. Acetate concentrations at each time point were calculated using an acetate standard curve generated in parallel. Initial velocities (*v*_0_) were determined from the linear portion of the reaction progress curves. Michaelis–Menten parameters (*K_M_* and *k_cat_*) were obtained by fitting initial velocity versus substrate concentration data to the Michaelis–Menten equation. Nonlinear regression analysis was performed in R.

#### In vitro deacetylase activity of immunopurified HDAC6

HDAC6-dependent AcK hydrolysis was assessed using immunoaffinity-purified protein from HEK293T cells. Briefly, 4.0 × 10^6^ HEK293T cells were seeded in T75 flasks and transfected the following day with 30 µg pCDNA3.1/HDAC6-Flag or empty pCDNA3.1 vector using PEI, and harvested 48 h post-transfection. Cell pellets were resuspended in 900 µL extraction buffer [20 mM HEPES (pH 7.9), 0.8 M NaCl, 0.1% Nonidet P-40, 25% glycerol] supplemented with DNase I and protease inhibitor cocktail and incubated on ice for 1 h. Lysates were clarified by centrifugation at 15,000×g for 10 min at 4*^◦^*C. Supernatants were incubated overnight at 4*^◦^*C with 50 µL anti-Flag magnetic beads with gentle rotation. Beads were washed three times with ice-cold PBS and twice with AcK deacetylation assay buffer [20 mM HEPES (pH 7.5), 100 mM NaCl, 5 mM KCl, 1 mM MgCl_2_], and resuspended in 200 µL of the same buffer. Initial in vitro deacetylation reactions were performed at 37*^◦^*C for 1 h. However, under these conditions, the H611A mutant retained ∼12% of WT activity. To improve enzyme stability in subsequent assays, 10% glycerol was included in all in vitro reaction buffers.

For LC–MS analysis, 0.5 mM AcK was added to 100 µL HDAC6-bound beads in a final volume of 120 µL LC–MS assay buffer [20 mM HEPES (pH 7.5), 100 mM NaCl, 5 mM KCl, 1 mM MgCl_2_, 0.2 mM phenylalanine, 10% glycerol]. Reactions were incubated at 37*^◦^*C, and 20 µL aliquots were withdrawn at 0, 2, 4, 6, and 16 h. Aliquots were quenched in an equal volume of 20% trichloroacetic acid (TCA) to precipitate proteins and centrifuged at 15,000×g for 30 min at 4*^◦^*C. Supernatants were diluted four-fold with triple-distilled water, and 25 µL was subjected to LC–MS analysis.

For plate reader–based quantification, 3 mM AcK was added to 32 µL HDAC6-bound beads in a final volume of 128 µL AcK deacetylation assay buffer supplemented with 10% glycerol. Reactions were incubated at 37*^◦^*C for 1 h. Aliquots (60 µL) were collected at 0 and 60 min, quenched with 2.5 µL 10% (v/v) HCl, neutralized with 15 µL freshly prepared 8% (w/v) sodium bicarbonate, and clarified by centrifugation at 12,000×g for 5 min. 35 µL of the clear supernatant was transferred to a 96-well microplate, and acetate production was quantified as described above.

#### In vitro deacetylase activity in whole-cell lysates

For lysate-based assays, HEK293T cells were transfected with vectors encoding WT or mutant HDAC6, or empty vector, using PEI. Cells were harvested 48 h post-transfection, washed with ice-cold PBS, flash-frozen in liquid nitrogen, and stored at-20*^◦^*C. Prior to analysis, cell pellets were thawed and resuspended in AcK deacetylation assay buffer supplemented with 10% glycerol, DNase I, and protease inhibitor cocktail, and incubated on ice for 20 min. Lysates were clarified by centrifugation at 15,000×g for 10 min at 4*^◦^*C. Total protein concentrations were determined using a BCA assay. AcK hydrolysis reactions were performed with 3 mM AcK and increasing concentrations of total lysate protein (0–2.5 µg/µL) at 37*^◦^*C for 1 h with gentle agitation. For comparison of HDAC6 variants, reactions contained 0.5 µg/µL total lysate protein and 3 mM AcK and were incubated for 1 h at 37*^◦^*C. Acetate production was quantified as described above. Relative enzymatic activity was calculated based on acetate generation normalized to the amount of HDAC6.

#### Fluor-de-Lys assay for HDAC6 CD2 activity

Deacetylase activity of HDAC6 and the indicated mutants toward a fluorogenic substrate was measured using a Fluor-de-Lys (FDL) assay based on an acetylated lysine–7-amino-4-methylcoumarin (AcK-AMC) substrate. Reactions were performed in FDL buffer [50 mM Tris-HCl (pH 8.0), 137 mM NaCl, 2.7 mM KCl, 1 mM MgCl_2_, 1 mg/mL BSA] in a microplate format. Recombinant HDAC6 WT or mutant proteins were used at a final concentration of 2 µM and incubated with 1.25 mM AcK-AMC substrate at 37*^◦^*C. Reactions were initiated by addition of substrate and carried out for the indicated time points. At each time point, reactions were terminated by addition of 54 µL developer solution (3 mg/mL trypsin, 0.05 mM HCl in FDL buffer) to allow proteolytic cleavage of deacetylated substrate. Samples were incubated at 25*^◦^*C for 1.5 h to ensure complete AMC release. Fluorescence was measured using a Spark microplate reader with excitation at 340 nm and emission at 465 nm. Background fluorescence from substrate-only controls was subtracted from all measurements. Enzymatic activity was calculated from the linear range of fluorescence increase over time and normalized to the amount of HDAC6.

#### In vitro SSAT2 acetyltransferase activity assay

SSAT2 activity was quantified using a colorimetric assay based on detection of free CoA generated during acetyl transfer from acetyl-CoA to lysine. Free CoA reacts with 5,5’-dithiobis-(2-nitrobenzoate) (DTNB) to produce 2-nitro-5-thiobenzoate, which absorbs at 412 nm. For assays using IPed SSAT2, reactions were performed in a total volume of 150 µL containing 150 mM Tris-HCl (pH 7.5), 2 mM EDTA, 5% glycerol, 1 mM acetyl-CoA, and 40 mM lysine. Following incubation at 30*^◦^*C for 4 h, 32 µL of the reaction mixture was transferred to a clear 96-well plate. DTNB (5 µL, 10 mM) and 63 µL 150 mM Tris-HCl (pH 7.5) were added to each well, and absorbance at 412 nm was measured using a plate reader. Activity was calculated after subtraction of blank values obtained from mock-transfected cell immunoprecipitates.

Measurements of purified recombinant SSAT2 (0.2 µM) were performed in 50 mM Bicine (pH 9.0) containing 1 mM acetyl-CoA and either 10 mM lysine or 10 mM thialysine. Reactions were incubated at 37*^◦^*C for 60 min and terminated by addition of 150 µL ethanol. Free CoA was detected by addition of 500 µL 0.2 mM DTNB prepared in 50 mM Tris-HCl (pH 8.0). After centrifugation at 15,000×g for 10 min at 4*^◦^*C, 100 µL of the supernatant was transferred to a 96-well plate, and absorbance at 412 nm was recorded. Control reactions lacking enzyme were used to determine background signal for each substrate. Enzymatic activity was calculated after blank subtraction.

#### In vitro autoacetylation assay

Autoacetylation of SSAT2 was assessed in vitro using purified recombinant proteins. Reactions were carried out in buffer containing 150 mM Tris-HCl (pH 8.5), 5% glycerol, and 10 mM MgCl_2_. Purified WT or R170 SSAT2 (25 µM) was incubated with 1.5 mM acetyl-CoA at 37*^◦^*C for 2 h. For auto-acetylation inhibition experiments, lysine (4 mM) was included in the reaction, and acetyl-CoA concentration was increased to 10 mM. Reactions were terminated by addition of 1× reducing Laemmli sample buffer. Samples were resolved by 14% SDS–PAGE and analyzed by immunoblotting using a pan–acetyl-lysine antibody.

### Protein electroporation

Recombinant protein delivery into cells was performed using the Neon Transfection System (Thermo Fisher) according to the manufacturer’s instructions with minor modifications. Purified SSAT2 and GFP-His control proteins were diluted to 50 µM in electroporation buffer [20 mM Tris-HCl (pH 8.0), 10% glycerol] immediately prior to use. WT and HDAC6-knockout HCT116 cells were washed twice with PBS and resuspended in Buffer R (Thermo Fisher) at a density of 4–8×10^7^ cells/mL. For each electroporation reaction, 4–8×10^5^ cells (12 µL) were mixed with 6 µL diluted SSAT2 and 6 µL diluted GFP-His protein (total volume 24 µL). Electroporation was performed at 1530 V, 20 ms, and 1 pulse using a 10 µL Neon tip. Immediately after electroporation, cells were transferred into pre-warmed, antibiotic-free McCoy’s 5A complete medium and cultured for 12 h. Cells were subsequently washed twice with PBS, flash-frozen in liquid nitrogen, and stored at-20*^◦^*C. For analysis, cells were lysed in RIPA buffer supplemented with protease inhibitor cocktail and clarified by centrifugation at 15,000×g for 10 min at 4*^◦^*C. Protein concentrations were determined using a BCA assay. Equal amounts of total protein (30 µg) were transferred to a black 96-well plate for quantification of GFP fluorescence using a plate reader, providing an internal normalization control. SSAT2 levels were analyzed by immunoblotting. Band intensities were quantified by densitometry and normalized to GFP fluorescence signals from the same samples. Data are presented relative to time zero as indicated in the figure legend.

### Western blotting analysis

Protein lysates and eluates were resolved on 10%, 12%, or 14% SDS–polyacrylamide gels supplemented with 1% TCE to enable rapid in-gel visualization of total protein. Following electrophoresis, gels were exposed to UV illumination to assess protein loading prior to membrane transfer. Proteins were then transferred onto 0.22 µm nitrocellulose membranes using a semi-dry transfer system (PowerBlotter Station, Invitrogen) according to the manufacturer’s instructions. Membranes were blocked for 1 h at room temperature in 5% (w/v) non-fat dry milk prepared in 1× TBST (Tris-buffered saline supplemented with 0.1% Tween-20), followed by incubation with the indicated primary antibodies overnight at 4*^◦^*C. Membranes were washed three times for 5 min each in 1× TBST and incubated with horseradish peroxidase (HRP)-conjugated secondary antibodies for 1 h at room temperature. After additional washes, immunoreactive bands were detected using enhanced chemiluminescence (ECL) reagent and imaged with a Fusion FX imaging system (Vilber). Band intensities were quantified by densitometry using ImageJ. ^77^

### In-gel trypsin digestion for NanoLC/MS

For analysis of site-specific acetyl-lysine incorporation, HEK293T cells (80% confluency; one 10 cm dish) were co-transfected with the SSAT2 expression plasmid and a vector encoding GFP-N150TAG-HA together with the AcK incorporation machinery. Forty-eight hours post-transfection, cells were harvested by scraping, pelleted by centrifugation at 2,000×g for 5 min at 4*^◦^*C, washed once with ice-cold PBS, flash-frozen in liquid nitrogen, and stored at-20*^◦^*C until further processing. For lysis, cell pellets were resuspended in nucleocytoplasmic extraction buffer [20 mM HEPES (pH 7.9), 0.8 M NaCl, 0.1% Nonidet P-40, 25% glycerol] supplemented with DNase I, 20 mM NAM, 6 µM SAHA, and protease inhibitor cocktail. Lysates were incubated on ice for 20 min and mechanically homogenized by passing 10 times through a 20-gauge needle. Insoluble material was removed by centrifugation at 15,000×g for 10 min at 4*^◦^*C. For immunoaffinity purification, 50 µL anti-HA magnetic beads were washed three times with extraction buffer and incubated with clarified lysates overnight at 4*^◦^*C with gentle rotation. Beads were subsequently washed five times with PBS to remove nonspecifically bound proteins. Bound proteins were eluted in 1× non-reducing Laemmli sample buffer (Bioprep, #LSB4-NR-5ml) at 70*^◦^*C for 10 min. Eluates were resolved by 12% Tris–glycine SDS–PAGE and visualized by staining with 0.1 (w/v) Coomassie Brilliant Blue R250. The band corresponding to GFP was excised, subjected to in-gel trypsin digestion following standard procedures, and analyzed by NanoLC–MS/MS as described previously.^37^. MS/MS spectra were searched against the appropriate protein database to confirm site-specific incorporation of AcK at the engineered position.

### Targeted metabolomics

Intracellular AcK levels were quantified by targeted LC–MS analysis performed at the Metabolomics Unit of Ben-Gurion University of the Negev. WT and HDAC6-knockout HCT116 cells (*n* ≥ 6) and HepG2 cells (*n* = 6) were cultured in 10 cm dishes to confluency. Cells were washed twice with 150 mM ammonium acetate to remove extracellular metabolites, flash-frozen in liquid nitrogen, and stored at-80*^◦^*C until extraction. Metabolites were extracted using methanol/water/chloroform (v/v = 2:1:1) as described previously,^78^ with modifications to cell culture conditions. Stable isotope-labeled internal standards (4.8 µg ^15^N-L-glutamine and 2.4 µg ^13^C_9_,^15^N-L-phenylalanine) were added at the beginning of extraction to enable normalization and control for technical variability. Extracts were dried completely in a vacuum concentrator and resuspended in 50 µL water/acetonitrile (v/v = 1:1) prior to analysis. Chromatographic separation was performed on a Waters Acquity UPLC system equipped with a BEH Amide HILIC column (1.7 µm, 2.1 × 100 mm) coupled to a Thermo Fisher Scientific Exploris 240 high-resolution mass spectrometer with electrospray ionization. The column temperature was maintained at 30*^◦^*C, the flow rate was 0.25 mL/min, and the injection volume was 1 µL. Mobile phase A consisted of 10 mM ammonium formate in 10% acetonitrile (pH 3.5), and mobile phase B consisted of 10 mM ammonium formate in 90% acetonitrile (pH 3.5). The gradient was as follows: 0–2 min, 5% A; 2–15 min, linear increase to 50% A; 15–17 min, 50% A; 17–18 min, linear increase to 70% A; 18–20 min, 70% A; 20–22 min, return to 5% A; 22–26 min, 5% A for re-equilibration. Data were acquired in positive ion full-scan mode (m/z 75–750) at a resolution of 240,000. MS source parameters were set according to the manufacturer’s recommendations for the selected flow rate. Peak extraction and integration were performed using Xcalibur QualBrowser (version 4.5.445). Target metabolites (AcK, lysine, ^15^N-L-glutamine, and ^13^C_9_,^15^N-L-phenylalanine) were identified by matching exact m/z values and retention times from extracted ion chromatograms to those of synthetic standards analyzed under identical conditions. Peak areas for AcK were first normalized to the internal standard signal and subsequently normalized to cell number. Relative metabolite abundances were compared across experimental groups as indicated.

LC-MS analysis of serum N-6-acetyl lysine was performed at the Metabolomic Center of Georgetown University. Blood was collected from age-matched HDAC6 knockout and wild-type C57Bl/6 mice (4-6 months) via the submandibular sampling method approved in the IACUC protocol (Protocol number: 2022-0016) at Georgetown University. About 100-120 µL of blood samples were collected in BD Microtainer® K2 EDTA Microtubes, and then centrifuged at 400×g for 10 min. The top layer of plasma was aliquoted and frozen at-80*^◦^*C. For LC-MS analysis, 125 µl of extraction buffer (methanol/water, v/v=1:1) containing ^15^N-L-tyrosine at a concentration of 1 µg/ml was added to 25 µL of plasma. The mixture was vortexed briefly, then incubated on ice for 20 min followed by incubation at-20*^◦^*C for another 20 min. Samples were centrifuged at 4*^◦^*C for 20 min at 13,000 rpm to pellet insoluble materials. The supernatant was transferred to LC-MS vials. 5 µL of the prepared sample was resolved on a CSH Phenyl-hexyl, 2.7 µm, 2.1×100 mm column online with a triple quadrupole mass spectrometer (Xevo-TQ-S, Waters Corporation, USA) operating in the multiple reaction monitoring (MRM) mode. The sample cone voltage and collision energies were optimized for the analyte to obtain maximum ion intensity for parent and daughter ions using the “IntelliStart” feature of MassLynx software (Waters Corporation, USA). Mobile phases were as follows: solvent A, water with 0.1% formic acid; solvent B, acetonitrile with 0.1% formic acid. The flow rate for each run started 2% B for 1 min at 0.3 mL/min, followed by a step gradient at 25% B for 1 min, followed by a linear gradient starting at 25% B changing to 95% B over the course of 2.6 min at 0.5 mL/min, followed by 5% B for 2 min at 0.3 mL/min. Data were processed using Target Lynx 4.1. Quantification of the endogenous metabolite concentrations was performed by generating an external standard curve with known concentrations of each metabolite. Metabolite standards were analyzed alongside the plasma samples using the same method in the same run. A calibration standard curve generated from the metabolite standard concentrations and total peak areas was used to calculate the concentrations of each endogenous metabolite.

### Statistics and reproducibility

Statistical analyses were performed using R (R Foundation for Statistical Computing). Unless otherwise indicated, comparisons among multiple groups were conducted using one-way analysis of variance (ANOVA) followed by Tukey’s post hoc test for multiple comparisons. A *p* value of ≤ 0.05 was considered statistically significant, and exact *p* values are provided in the figures. Data are presented as mean ± standard deviation (SD) unless otherwise indicated. Unless otherwise indicated, all experiments were independently repeated at least three times with comparable results, and representative data are shown where applicable. Investigators were not blinded to group allocation during experiments or during outcome assessment, except for the transwell cell migration assays, in which investigators were blinded to sample identity.

## Acknowledgements

This work was supported by the European Research Council (ERC) under the European Union Horizon 2020 research and innovation programme [grant number 678461 to E.A.]; the Israel Science Foundation [grants number 1769/20 and 2331/25 to E.A.]; and the Israeli Ministry of Innovation, Science, and Technology [grant number 5936 to E.A.]. This work was in part supported by the CAS (RVO: 86652036 to C.B.).

## Author contributions

Conceptualization: F.W., and E.A. Investigation: F.W. conducted most of the experiments; S.P. contributed to in vitro zCD1 activity measurements; S.N. performed and analyzed the metabolomic measurements of mouse models; A.B. performed and analyzed the metabolomic measurements of cells; M.V. contributed to in vitro HDAC6 activity measurements. Formal analysis: F.W. and E.A. Visualization: F.W. and E.A. Funding acquisition: C.B. and E.A. Supervision: A.V., C.B., and E.A. Writing: F.W. and E.A. All authors commented on the manuscript.

## REFERENCES

1. Choudhary, C., Kumar, C., Gnad, F., Nielsen, M.L., Rehman, M., Walther, T.C., Olsen, J.V., and Mann, M. (2009). Lysine Acetylation Targets Protein Complexes and Co-Regulates Major Cellular Functions. Science 325, 834–840. 10.1126/science.1175371.

2. Carrico, C., Meyer, J.G., He, W., Gibson, B.W., and Verdin, E. (2018). The Mitochondrial Acylome Emerges: Proteomics, Regulation by Sirtuins, and Metabolic and Disease Implications. Cell Metabolism 27, 497–512. 10.1016/j.cmet.2018.01.016.

3. Ali, I., Conrad, R.J., Verdin, E., and Ott, M. (2018). Lysine Acetylation Goes Global: From Epigenetics to Metabolism and Therapeutics. Chemical Reviews 118, 1216–1252. 10.1021/acs.chemrev.7b00181.

4. Choudhary, C., Weinert, B.T., Nishida, Y., Verdin, E., and Mann, M. (2014). The Growing Landscape of Lysine Acetylation Links Metabolism and Cell Signalling. Nature Reviews. Molecular Cell Biology 15, 536–550. 10.1038/nrm3841.

5. Shvedunova, M. and Akhtar, A. (2022). Modulation of Cellular Processes by Histone and Non-Histone Protein Acetylation. Nature Reviews. Molecular Cell Biology 23, 329–349. 10.1038/s41580-021-00441-y.

6. Chaleckis, R., Ebe, M., Pluskal, T., Murakami, I., Kondoh, H., and Yanagida, M. (2014). Unexpected Similarities between the Schizosaccharomyces and Human Blood Metabolomes, and Novel Human Metabolites. Molecular bioSystems 10, 2538–2551. 10.1039/c4mb00346b.

7. Chaleckis, R., Murakami, I., Takada, J., Kondoh, H., and Yanagida, M. (2016). Individual Variability in Human Blood Metabolites Identifies Age-Related Differences. Proceedings of the National Academy of Sciences of the United States of America 113, 4252–4259. 10.1073/pnas.1603023113.

8. Huang, T., Zeleznik, O.A., Poole, E.M., Clish, C.B., Deik, A.A., Scott, J.M., Vetter, C., Schernhammer, E.S., Brunner, R., Hale, L., Manson, J.E., Hu, F.B., Redline, S., Tworoger, S.S., and Rexrode, K.M. (2019). Habitual Sleep Quality, Plasma Metabolites and Risk of Coronary Heart Disease in Post-Menopausal Women. International Journal of Epidemiology 48, 1262–1274. 10.1093/ije/dyy234.

9. Jalaleddine, N., Hachim, M., Al-Hroub, H., Saheb Sharif-Askari, N., Senok, A., Elmoselhi, A., Mahboub, B., Samuel Kurien, N.M., Kandasamy, R.K., Semreen, M.H., Halwani, R., Soares, N.C., and Al Heialy, S. (2022). N6-Acetyl-L-Lysine and p-Cresol as Key Metabolites in the Pathogenesis of COVID-19 in Obese Patients. Frontiers in Immunology 13, 827603. 10.3389/fimmu.2022.827603.

10. Yin, X., Chan, L.S., Bose, D., Jackson, A.U., VandeHaar, P., Locke, A.E., Fuchsberger, C., Stringham, H.M., Welch, R., Yu, K., Fernandes Silva, L., Service, S.K., Zhang, D., Hector, E.C., Young, E., Ganel, L., Das, I., Abel, H., Erdos, M.R., Bonnycastle, L.L., Kuusisto, J., Stitziel, N.O., Hall, I.M., Wagner, G.R., FinnGen, Kang, J., Morrison, J., Burant, C.F., Collins, F.S., Ripatti, S., Palotie, A., Freimer, N.B., Mohlke, K.L., Scott, L.J., Wen, X., Fauman, E.B., Laakso, M., and Boehnke, M. (2022). Genome-Wide Association Studies of Metabolites in Finnish Men Identify Disease-Relevant Loci. Nature Communications 13, 1644. 10.1038/s41467-022-29143-5.

11. Feofanova, E.V., Brown, M.R., Alkis, T., Manuel, A.M., Li, X., Tahir, U.A., Li, Z., Mendez, K.M., Kelly, R.S., Qi, Q., Chen, H., Larson, M.G., Lemaitre, R.N., Morrison, A.C., Grieser, C., Wong, K.E., Gerszten, R.E., Zhao, Z., Lasky-Su, J., NHLBI Trans-Omics for Precision Medicine (TOPMed), and Yu, B. (2023). Whole-Genome Sequencing Analysis of Human Metabolome in Multi-Ethnic Populations. Nature Communications 14, 3111. 10.1038/s41467-023-38800-2.

12. Dong, T., Wang, X., Jia, Z., Yang, J., and Liu, Y. (2024). Assessing the Associations of 1,400 Blood Metabolites with Major Depressive Disorder: A Mendelian Randomization Study. Frontiers in Psychiatry 15, 1391535. 10.3389/fpsyt.2024.1391535.

13. Rinschen, M.M., Palygin, O., El-Meanawy, A., Domingo-Almenara, X., Palermo, A., Dissanayake, L.V., Golosova, D., Schafroth, M.A., Guijas, C., Demir, F., Jaegers, J., Gliozzi, M.L., Xue, J., Hoehne, M., Benzing, T., Kok, B.P., Saez, E., Bleich, M., Himmerkus, N., Weisz, O.A., Cravatt, B.F., Krüger, M., Benton, H.P., Siuzdak, G., and Staruschenko, A. (2022). Accelerated Lysine Metabolism Conveys Kidney Protection in Salt-Sensitive Hypertension. Nature Communications 13, 4099. 10.1038/s41467-022-31670-0.

14. Hunsberger, H.C., Greenwood, B.P., Tolstikov, V., Narain, N.R., Kiebish, M.A., and Denny, C.A. (2020). Divergence in the Metabolome between Natural Aging and Alzheimer’s Disease. Scientific Reports 10, 12171. 10.1038/s41598-020-68739-z.

15. Guo, D., Li, N., Zhang, X., Zhou, R., He, J., Ding, X.P., Yu, W., Tong, F., Yin, S., Wang, Y., Xu, X., Wang, L., Fan, M., Feng, S., Liu, K., Tang, K., Ouyang, Z., Guo, Y.R., and Wang, Y. (2024). Co-Translational Deposition of N6-Acetyl-L-Lysine in Nascent Proteins Contributes to the Acetylome in Mammalian Cells. Advanced Science (Weinheim, Baden-Wurttemberg, Germany) e2403309. 10.1002/advs.202403309.

16. Rajantie, J., Simell, O., and Perheentupa, J. (1983). Oral Administration of Epsilon N-acetyllysine and Homocitrulline in Lysinuric Protein Intolerance. Journal of Pediatrics 102, 388–390. 10.1016/s0022-3476(83)80654-4.

17. Wang, Z., Zang, C., Cui, K., Schones, D.E., Barski, A., Peng, W., and Zhao, K. (2009). Genome-Wide Mapping of HATs and HDACs Reveals Distinct Functions in Active and Inactive Genes. Cell 138, 1019–1031. 10.1016/j.cell.2009.06.049.

18. Gregoretti, I.V., Lee, Y.M., and Goodson, H.V. (2004). Molecular Evolution of the Histone Deacetylase Family: Functional Implications of Phylogenetic Analysis. Journal of Molecular Biology 338, 17–31. 10.1016/j.jmb.2004.02.006.

19. Roth, S.Y., Denu, J.M., and Allis, C.D. (2001). Histone Acetyltransferases. Annual Review of Biochemistry 70, 81–120. 10.1146/annurev.biochem.70.1.81.

20. Paik, W.K., Bloch-Frankenthal, L., Birnbaum, S.M., Winitz, M., and Greenstein, J.P. (1957). Epsilon-Lysine Acylase. Archives of Biochemistry and Biophysics 69, 56–66. 10.1016/0003-9861(57)90472-1.

21. Hysi, P.G., Mangino, M., Christofidou, P., Falchi, M., Karoly, E.D., Nihr Bioresource Investigators, n., Mohney, R.P., Valdes, A.M., Spector, T.D., and Menni, C. (2022). Metabolome Genome-Wide Association Study Identifies 74 Novel Genomic Regions Influencing Plasma Metabolites Levels. Metabolites 12, 61. 10.3390/metabo12010061.

22. Chen, Y., Lu, T., Pettersson-Kymmer, U., Stewart, I.D., Butler-Laporte, G., Nakanishi, T., Cerani, A., Liang, K.Y.H., Yoshiji, S., Willett, J.D.S., Su, C.Y., Raina, P., Greenwood, C.M.T., Farjoun, Y., Forgetta, V., Langenberg, C., Zhou, S., Ohlsson, C., and Richards, J.B. (2023). Genomic Atlas of the Plasma Metabolome Prioritizes Metabolites Implicated in Human Diseases. Nature Genetics 55, 44–53. 10.1038/s41588-022-01270-1.

23. Hai, Y. and Christianson, D.W. (2016). Histone Deacetylase 6 Structure and Molecular Basis of Catalysis and Inhibition. Nature Chemical Biology 12, 741–747. 10.1038/nchembio.2134.

24. Miyake, Y., Keusch, J.J., Wang, L., Saito, M., Hess, D., Wang, X., Melancon, B.J., Helquist, P., Gut, H., and Matthias, P. (2016). Structural Insights into HDAC6 Tubulin Deacetylation and Its Selective Inhibition. Nature Chemical Biology 12, 748–754. 10.1038/nchembio.2140.

25. Kovacs, J.J., Murphy, P.J., Gaillard, S., Zhao, X., Wu, J.T., Nicchitta, C.V., Yoshida, M., Toft, D.O., Pratt, W.B., and Yao, T.P. (2005). HDAC6 Regulates Hsp90 Acetylation and Chaperone-Dependent Activation of Glucocorticoid Receptor. Molecular Cell 18, 601–607. 10.1016/j.molcel.2005.04.021.

26. Hubbert, C., Guardiola, A., Shao, R., Kawaguchi, Y., Ito, A., Nixon, A., Yoshida, M., Wang, X.F., and Yao, T.P. (2002). HDAC6 Is a Microtubule-Associated Deacetylase. Nature 417, 455–458. 10.1038/417455a.

27. Haggarty, S.J., Koeller, K.M., Wong, J.C., Grozinger, C.M., and Schreiber, S.L. (2003). Domain-Selective Small-Molecule Inhibitor of Histone Deacetylase 6 (HDAC6)-Mediated Tubulin Deacetylation. Proceedings of the National Academy of Sciences of the United States of America 100, 4389–4394. 10.1073/pnas.0430973100.

28. Zhang, X., Yuan, Z., Zhang, Y., Yong, S., Salas-Burgos, A., Koomen, J., Olashaw, N., Parsons, J.T., Yang, X.J., Dent, S.R., Yao, T.P., Lane, W.S., and Seto, E. (2007). HDAC6 Modulates Cell Motility by Altering the Acetylation Level of Cortactin. Molecular Cell 27, 197–213. 10.1016/j.molcel.2007.05.033.

29. Kutil, Z., Skultetyova, L., Rauh, D., Meleshin, M., Snajdr, I., Novakova, Z., Mikesova, J., Pavlicek, J., Hadzima, M., Baranova, P., Havlinova, B., Majer, P., Schutkowski, M., and Barinka, C. (2019). The Unraveling of Substrate Specificity of Histone Deacetylase 6 Domains Using Acetylome Peptide Microarrays and Peptide Libraries. FASEB Journal 33, 4035–4045. 10.1096/fj.201801680R.

30. Chen, Y., Vujcic, S., Liang, P., Diegelman, P., Kramer, D.L., and Porter, C.W. (2003). Genomic Identification and Biochemical Characterization of a Second Spermidine/Spermine N1-acetyltransferase. Biochemical Journal 373, 661–667. 10.1042/BJ20030734.

31. Abo-Dalo, B., Ndjonka, D., Pinnen, F., Liebau, E., and Lüersen, K. (2004). A Novel Member of the GCN5-related N-acetyltransferase Superfamily from Caenorhabditis Elegans Preferentially Catalyses the N-acetylation of Thialysine [S-(2-Aminoethyl)-L-cysteine]. Biochemical Journal 384, 129–137. 10.1042/BJ20040789.

32. Coleman, C.S., Stanley, B.A., Jones, A.D., and Pegg, A.E. (2004). Spermidine/Spermine-N1-acetyltransferase-2 (SSAT2) Acetylates Thialysine and Is Not Involved in Polyamine Metabolism. The Biochemical Journal 384, 139–148. 10.1042/BJ20040790.

33. Lüersen, K. (2005). Leishmania Major Thialysine Nepsilon-acetyltransferase: Identification of Amino Acid Residues Crucial for Substrate Binding. FEBS Letters 579, 5347–5352. 10.1016/j.febslet.2005.08.063.

34. Bělíček, J., Ľuptáková, E., Kopečný, D., Frömmel, J., Vigouroux, A., Ćavar Zeljković, S., Jagic, F., Briozzo, P., Kopečný, D.J., Tarkowski, P., Nisler, J., De Diego, N., Moréra, S., and Kopečná, M. (2023). Biochemical and Structural Basis of Polyamine, Lysine and Ornithine Acetylation Catalyzed by Spermine/Spermidine N-acetyl Transferase in Moss and Maize. The Plant Journal: For Cell and Molecular Biology 114, 482–498. 10.1111/tpj.16148.

35. Vogel, N.L., Boeke, M., and Ashburner, B.P. (2006). Spermidine/Spermine N1-Acetyltransferase 2 (SSAT2) Functions as a Coactivator for NF-kappaB and Cooperates with CBP and P/CAF to Enhance NF-kappaB-dependent Transcription. Biochimica et Biophysica Acta 1759, 470–477. 10.1016/j.bbaexp.2006.08.005.

36. Baek, J.H., Liu, Y.V., McDonald, K.R., Wesley, J.B., Hubbi, M.E., Byun, H., and Semenza, G.L. (2007). Spermidine/Spermine-N1-acetyltransferase 2 Is an Essential Component of the Ubiquitin Ligase Complex That Regulates Hypoxia-Inducible Factor 1alpha. Journal of Biological Chemistry 282, 23572–23580. 10.1074/jbc.M703504200.

37. Wu, F., Muskat, N.H., Dvilansky, I., Koren, O., Shahar, A., Gazit, R., Elia, N., and Arbely, E. (2023). Acetylation-Dependent Coupling between G6PD Activity and Apoptotic Signaling. Nature Communications 14, 6208. 10.1038/s41467-023-41895-2.

38. Passaro, S., Corso, G., Wohlwend, J., Reveiz, M., Thaler, S., Somnath, V.R., Getz, N., Portnoi, T., Roy, J., Stark, H., Kwabi-Addo, D., Beaini, D., Jaakkola, T., and Barzilay, R. (2025). Boltz-2: Towards Accurate and Efficient Binding Affinity Prediction. Biorxiv: the Preprint Server for Biology 2025.6.14.659707. 10.1101/2025.06.14.659707.

39. Zou, H., Wu, Y., Navre, M., and Sang, B.C. (2006). Characterization of the Two Catalytic Domains in Histone Deacetylase 6. Biochemical and Biophysical Research Communications 341, 45–50. 10.1016/j.bbrc.2005.12.144.

40. Shukla, S., Komarek, J., Novakova, Z., Nedvedova, J., Ustinova, K., Vankova, P., Kadek, A., Uetrecht, C., Mertens, H., and Barinka, C. (2023). In-Solution Structure and Oligomerization of Human Histone Deacetylase 6 - an Integrative Approach. The FEBS journal 290, 821–836. 10.1111/febs.16616.

41. Zhang, Y., Gilquin, B., Khochbin, S., and Matthias, P. (2006). Two Catalytic Domains Are Required for Protein Deacetylation. The Journal of Biological Chemistry 281, 2401–2404. 10.1074/jbc.C500241200.

42. Ustinova, K., Novakova, Z., Saito, M., Meleshin, M., Mikesova, J., Kutil, Z., Baranova, P., Havlinova, B., Schutkowski, M., Matthias, P., and Barinka, C. (2020). The Disordered N-terminus of HDAC6 Is a Microtubule-Binding Domain Critical for Efficient Tubulin Deacetylation. Journal of Biological Chemistry 295, 2614–2628. 10.1074/jbc.RA119.011243.

43. Langousis, G., Sanchez, J., Kempf, G., and Matthias, P. (2023). Expression and Crystallization of HDAC6 Tandem Catalytic Domains. Methods in Molecular Biology (clifton, N.J.) 2589, 467–480. 10.1007/978-1-0716-2788-4_30.

44. Donker, L. and Godinho, S.A. (2025). Rethinking Tubulin Acetylation: From Regulation to Cellular Adaptation. Current Opinion in Cell Biology 94, 102512. 10.1016/j.ceb.2025.102512.

45. Portran, D., Schaedel, L., Xu, Z., Théry, M., and Nachury, M.V. (2017). Tubulin Acetylation Protects Long-Lived Microtubules against Mechanical Ageing. Nature Cell Biology 19, 391–398. 10.1038/ncb3481.

46. Perdiz, D., Mackeh, R., Poüs, C., and Baillet, A. (2011). The Ins and Outs of Tubulin Acetylation: More than Just a Post-Translational Modification? Cellular Signalling 23, 763–771. 10.1016/j.cellsig.2010.10.014.

47. Balmer, M.L., Ma, E.H., Bantug, G.R., Grählert, J., Pfister, S., Glatter, T., Jauch, A., Dimeloe, S., Slack, E., Dehio, P., Krzyzaniak, M.A., King, C.G., Burgener, A.V., Fischer, M., Develioglu, L., Belle, R., Recher, M., Bonilla, W.V., Macpherson, A.J., Hapfelmeier, S., Jones, R.G., and Hess, C. (2016). Memory CD8(+) T Cells Require Increased Concentrations of Acetate Induced by Stress for Optimal Function. Immunity 44, 1312–1324. 10.1016/j.immuni.2016.03.016.

48. Schug, Z.T., Peck, B., Jones, D.T., Zhang, Q., Grosskurth, S., Alam, I.S., Goodwin, L.M., Smethurst, E., Mason, S., Blyth, K., McGarry, L., James, D., Shanks, E., Kalna, G., Saunders, R.E., Jiang, M., Howell, M., Lassailly, F., Thin, M.Z., Spencer-Dene, B., Stamp, G., van den Broek, N.J.F., Mackay, G., Bulusu, V., Kamphorst, J.J., Tardito, S., Strachan, D., Harris, A.L., Aboagye, E.O., Critchlow, S.E., Wakelam, M.J.O., Schulze, A., and Gottlieb, E. (2015). Acetyl-CoA Synthetase 2 Promotes Acetate Utilization and Maintains Cancer Cell Growth under Metabolic Stress. Cancer Cell 27, 57–71. 10.1016/j.ccell.2014.12.002.

49. Chen, S., Francioli, L.C., Goodrich, J.K., Collins, R.L., Kanai, M., Wang, Q., Alföldi, J., Watts, N.A., Vittal, C., Gauthier, L.D., et al. (2024). A Genomic Mutational Constraint Map Using Variation in 76,156 Human Genomes. Nature 625, 92–100. 10.1038/s41586-023-06045-0.

50. Akimov, V., Barrio-Hernandez, I., Hansen, S.V.F., Hallenborg, P., Pedersen, A.K., Bekker-Jensen, D.B., Puglia, M., Christensen, S.D.K., Vanselow, J.T., Nielsen, M.M., Kratchmarova, I., Kelstrup, C.D., Olsen, J.V., and Blagoev, B. (2018). UbiSite Approach for Comprehensive Mapping of Lysine and N-terminal Ubiquitination Sites. Nature Structural & Molecular Biology 25, 631–640. 10.1038/s41594-018-0084-y.

51. Bewley, M.C., Graziano, V., Jiang, J., Matz, E., Studier, F.W., Pegg, A.E., Coleman, C.S., and Flanagan, J.M. (2006). Structures of Wild-Type and Mutant Human Spermidine/Spermine N1-acetyltransferase, a Potential Therapeutic Drug Target. Proceedings of the National Academy of Sciences of the United States of America 103, 2063–2068. 10.1073/pnas.0511008103.

52. Cai, S., Liu, X., Zhang, C., Xing, B., and Du, X. (2017). Autoacetylation of NAT10 Is Critical for Its Function in rRNA Transcription Activation. Biochemical and Biophysical Research Communications 483, 624–629. 10.1016/j.bbrc.2016.12.092.

53. Majorek, K.A., Kuhn, M.L., Chruszcz, M., Anderson, W.F., and Minor, W. (2013). Structural, Functional, and Inhibition Studies of a Gcn5-related N-acetyltransferase (GNAT) Superfamily Protein PA4794: A New C-terminal Lysine Protein Acetyltransferase from Pseudomonas Aeruginosa. Journal of Biological Chemistry 288, 30223–30235. 10.1074/jbc.M113.501353.

54. Liang, X., Xiao, J., Li, X., Liu, Y., Lu, Y., Wen, Y., Li, Z., Che, X., Ma, Y., Zhang, X., Zhang, Y., Jian, D., Wang, P., Xuan, C., Yu, G., Li, L., and Zhang, H. (2022). A C-terminal Glutamine Recognition Mechanism Revealed by E3 Ligase TRIM7 Structures. Nature Chemical Biology 18, 1214–1223. 10.1038/s41589-022-01128-x.

55. Ru, Y., Yan, X., Zhang, B., Song, L., Feng, Q., Ye, C., Zhou, Z., Yang, Z., Li, Y., Zhang, Z., Li, Q., Mi, W., and Dong, C. (2022). C-Terminal Glutamine Acts as a C-degron Targeted by E3 Ubiquitin Ligase TRIM7. Proceedings of the National Academy of Sciences of the United States of America 119, e2203218119. 10.1073/pnas.2203218119.

56. Baker, S.A. and Rutter, J. (2023). Metabolites as Signalling Molecules. Nature Reviews. Molecular Cell Biology 24, 355–374. 10.1038/s41580-022-00572-w.

57. Zhang, B. and Schroeder, F.C. (2025). Mechanisms of Metabolism-Coupled Protein Modifications. Nature Chemical Biology 21, 819–830. 10.1038/s41589-024-01805-z.

58. Hai, Y., Shinsky, S.A., Porter, N.J., and Christianson, D.W. (2017). Histone Deacetylase 10 Structure and Molecular Function as a Polyamine Deacetylase. Nature Communications 8, 15368. 10.1038/ncomms15368.

59. Hegde, S.S., Chandler, J., Vetting, M.W., Yu, M., and Blanchard, J.S. (2007). Mechanistic and Structural Analysis of Human Spermidine/Spermine N1-acetyltransferase. Biochemistry 46, 7187–7195. 10.1021/bi700256z.

60. Cao, Y., Zhao, Y., Deng, T., Zhou, Q., Hu, G., Hu, Z.L., Jiang, Y.Y., Yang, X.H., Wang, F., Wu, P.F., and Chen, J.G. (2025). Hepatic Acetyl-CoA Metabolism Modulates Neuroinflammation and Depression Susceptibility via Acetate. Cell Metabolism 37, 2185–2201.e8. 10.1016/j.cmet.2025.08.010.

61. Klunk, W.E., McClure, R.J., and Pettegrew, J.W. (1991). L-Phosphoserine, a Metabolite Elevated in Alzheimer’s Disease, Interacts with Specific L-glutamate Receptor Subtypes. Journal of Neurochemistry 56, 1997–2003. 10.1111/j.1471-4159.1991.tb03458.x.

62. Hoppel, C.L., Cox, R.A., and Novak, R.F. (1980). N6-Trimethyl-lysine Metabolism. 3-Hydroxy-N6-trimethyl-lysine and Carnitine Biosynthesis. Biochemical Journal 188, 509–519. 10.1042/bj1880509.

63. Matthews, D.E. (2020). Review of Lysine Metabolism with a Focus on Humans. The Journal of Nutrition 150, 2548S–2555S. 10.1093/jn/nxaa224.

64. Yamashita, K. and Ashida, K. (1969). Lysine Metabolism in Rats Fed Lysine-free Diet. Journal of Nutrition 99, 267–273. 10.1093/jn/99.3.267.

65. Said, A.K., Hegsted, D.M., and Hayes, K.C. (1974). Response of Adult Rats to Deficiencies of Different Essential Amino Acids. British Journal of Nutrition 31, 47–57. 10.1079/BJN19740007.

66. Jochems, J., Boulden, J., Lee, B.G., Blendy, J.A., Jarpe, M., Mazitschek, R., Van Duzer, J.H., Jones, S., and Berton, O. (2014). Antidepressant-like Properties of Novel HDAC6-selective Inhibitors with Improved Brain Bioavailability. Neuropsychopharmacology: Official Publication of the American College of Neuropsychopharmacology 39, 389–400. 10.1038/npp.2013.207.

67. Meylan, E.M., Halfon, O., Magistretti, P.J., and Cardinaux, J.R. (2016). The HDAC Inhibitor SAHA Improves Depressive-like Behavior of CRTC1-deficient Mice: Possible Relevance for Treatment-Resistant Depression. Neuropharmacology 107, 111–121. 10.1016/j.neuropharm.2016.03.012.

68. Fukada, M., Hanai, A., Nakayama, A., Suzuki, T., Miyata, N., Rodriguiz, R.M., Wetsel, W.C., Yao, T.P., and Kawaguchi, Y. (2012). Loss of Deacetylation Activity of Hdac6 Affects Emotional Behavior in Mice. PloS One 7, e30924. 10.1371/journal.pone.0030924.

69. Matsuzaki, J., Kurokawa, S., Iwamoto, C., Miyaho, K., Takamiya, A., Ishii, C., Hirayama, A., Sanada, K., Fukuda, S., Mimura, M., Kishimoto, T., and Saito, Y. (2024). Intestinal Metabolites Predict Treatment Resistance of Patients with Depression and Anxiety. Gut Pathogens 16, 8. 10.1186/s13099-024-00601-3.

70. Ryu, H.W., Won, H.R., Lee, D.H., and Kwon, S.H. (2017). HDAC6 Regulates Sensitivity to Cell Death in Response to Stress and Post-Stress Recovery. Cell Stress & Chaperones 22, 253–261. 10.1007/s12192-017-0763-3.

71. Avrahami, E.M., Levi, S., Zajfman, E., Regev, C., Ben-David, O., and Arbely, E. (2018). Reconstitution of Mammalian Enzymatic Deacylation Reactions in Live Bacteria Using Native Acylated Substrates. ACS Synthetic Biology 7, 2348–2354. 10.1021/acssynbio.8b00314.

72. Neumann, H., Hancock, S.M., Buning, R., Routh, A., Chapman, L., Somers, J., Owen-Hughes, T., van Noort, J., Rhodes, D., and Chin, J.W. (2009). A Method for Genetically Installing Site-Specific Acetylation in Recombinant Histones Defines the Effects of H3 K56 Acetylation. Molecular Cell 36, 153–163. 10.1016/j.molcel.2009.07.027.

73. Cohen, S. and Arbely, E. (2016). Single-Plasmid-Based System for Efficient Noncanonical Amino Acid Mutagenesis in Cultured Mammalian Cells. ChemBioChem 17, 1008–1011. 10.1002/cbic.201500681.

74. Aloush, N., Schvartz, T., König, A.I., Cohen, S., Brozgol, E., Tam, B., Nachmias, D., Ben-David, O., Garini, Y., Elia, N., and Arbely, E. (2018). Live Cell Imaging of Bioorthogonally Labelled Proteins Generated With a Single Pyrrolysine tRNA Gene. Scientific Reports 8, 14527. 10.1038/s41598-018-32824-1.

75. McNeal, T.A., Weinberger, J., Liman, G.L.S., Ariagno, T.M., Wood, D.W., Santangelo, T.J., and Lennon, C.W. (2025). Controllable Intein Splicing and N-terminal Cleavage at Mesophilic Temperatures. Frontiers in Bioengineering and Biotechnology 13, 1543573. 10.3389/fbioe.2025.1543573.

76. Qiu, L., Xu, W., Lu, X., Chen, F., Chen, Y., Tian, Y., Zhu, Q., Liu, X., Wang, Y., Pei, X.H., Xu, X., Zhang, J., and Zhu, W.G. (2023). The HDAC6-RNF168 Axis Regulates H2A/H2A.X Ubiquitination to Enable Double-Strand Break Repair. Nucleic Acids Research 51, 9166–9182. 10.1093/nar/gkad631.

77. Schneider, C.A., Rasband, W.S., and Eliceiri, K.W. (2012). NIH Image to ImageJ: 25 Years of Image Analysis. Nature Methods 9, 671–675. 10.1038/nmeth.2089.

78. Batushansky, A., Matsuzaki, S., Newhardt, M.F., West, M.S., Griffin, T.M., and Humphries, K.M. (2019). GC-MS Metabolic Profiling Reveals Fructose-2,6-Bisphosphate Regulates Branched Chain Amino Acid Metabolism in the Heart during Fasting. Metabolomics: Official Journal of the Metabolomic Society 15, 18. 10.1007/s11306-019-1478-5.

