## Supplementary Information for "HDAC6 and SSAT2 orchestrate acetyllysine metabolism and protein homeostasis"

### Supplementary Figures

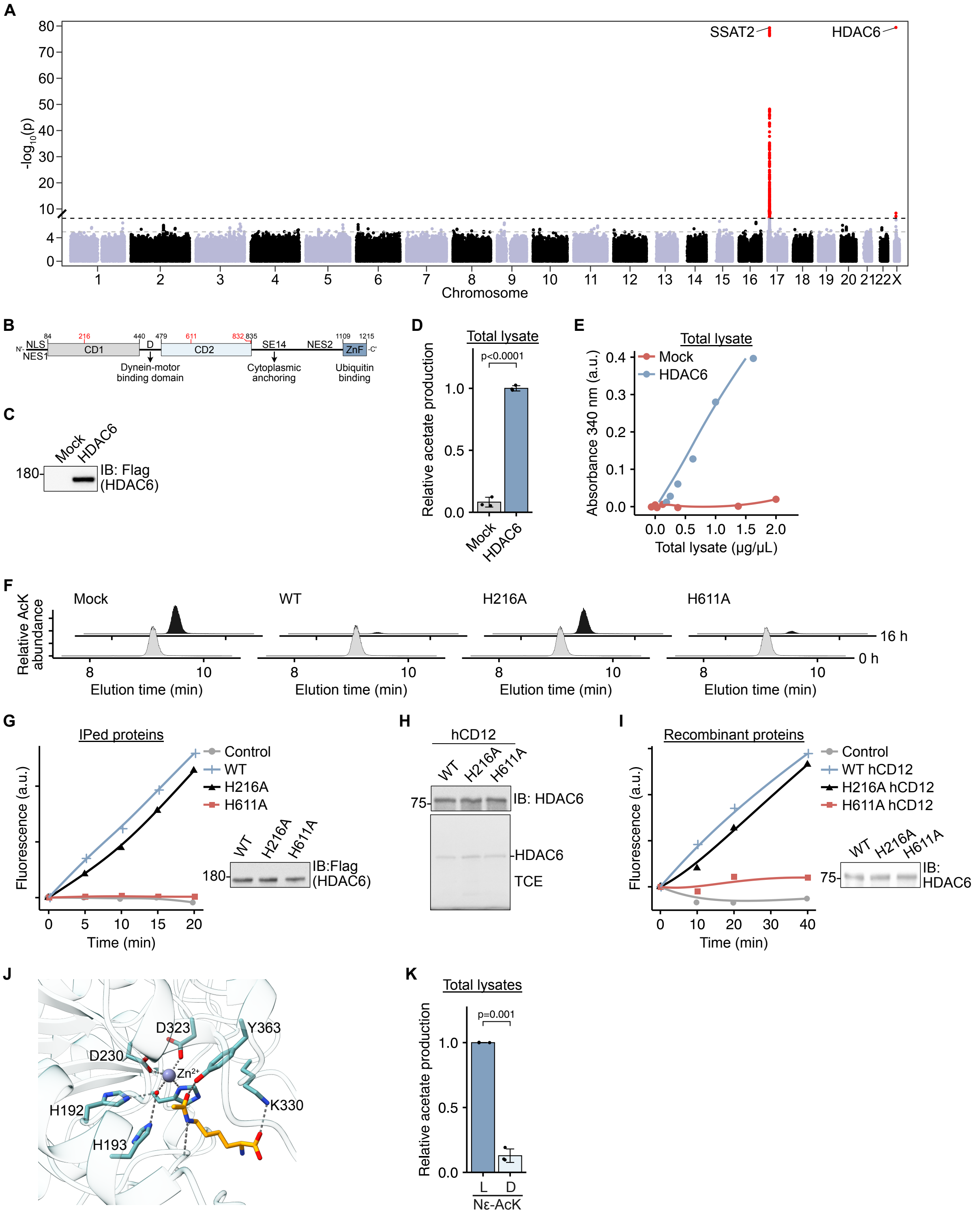


**Figure S1**

**Figure S1. Additional characterization of HDAC6 CD1 activity in vitro, related to Figure 1.**
(A) PheWeb plot of significant associations between SNPs across the human genome and abnormal AcK levels from the METSIM study GWAS.[1] Statistically significant associations ($p < 5 \times 10^{-8}$) are highlighted in red. Notably, variants within or near SSAT2 and HDAC6 show strong associations.
(B) Domain organization of HDAC6. HDAC6 is 1215 amino acids long and composed of five domains: N-terminus (residues 1–83), CD1 (residues 84–440), CD2 (residues 479–835), serine/glutamate-rich repeat motifs (residues 856–1108), and a C-terminal ZnF ubiquitin-binding domain (residues 1109–1215).
(C) Immunoblot using anti-Flag antibody showing the expression of HDAC6-Flag in HEK293T cells transfected with either HDAC6-Flag-expressing plasmid or empty vector.
(D) Relative acetate production following incubation of 3 mM AcK with 0.5 µg/µL total protein in cleared lysates of HEK293T cells transfected with either HDAC6-Flag-expressing plasmid or empty vector.
(E) Acetate production, measured by absorbance at 340 nm, following incubation of 3 mM AcK with cleared lysates of HEK293T cells transfected with either HDAC6-Flag-expressing plasmid or empty vector. Data are presented as a function of total protein in cleared lysates. Lines serve as guides to the eye.
(F) LC-MS analysis showing the relative abundance of AcK following incubation of 0.5 mM AcK with Flag-isolated proteins from cleared lysates of HEK293T cells transfected with either indicated HDAC6-Flag-expressing plasmid or empty vector.
(G) In vitro CD2 deacetylase activity of immunopurified WT or mutant Flag-HDAC6 described in panel F, using the commercially available fluorogenic Fluor-de-lys assay. Fluorescence, measured following substrate deacetylation, is plotted as a function of incubation time. Lines serve as guides for the eye. The immunoblot using anti-Flag antibody shows the total amount of each immunopurified protein used in the assay.
(H) Western blot analysis of truncated HDAC6 version that includes CD1 and CD2 (aa 84–835; hCD12). The WT and indicated mutants of hCD12 were recombinantly expressed in *E. coli*, purified, and probed with anti-HDAC6 antibody.
(I) In vitro CD2 deacetylase activity of purified WT or mutant hCD12 described in panel H. Deacetylase activity was measured as described in panel G. The immunoblot using anti-HDAC6 antibody shows the total amount of each protein used in the assay.
(J) Model of AcK bound in the active site of zCD1. Electrostatic interactions, including the interaction between the negatively charged Cα carboxylic group and positively charged K330, are represented as a dashed line.
(K) Relative acetate production following incubation of Nε-L- or D-acetyllysine with cleared lysates of HEK293T cells expressing HDAC6-Flag.
Data represent the mean ± S.D. of at least three independent experiments. Statistical analysis was performed using the Welch two-sample t-test (D, K). ns, non-significant. In-gel TCE fluorescence serves as a loading control.

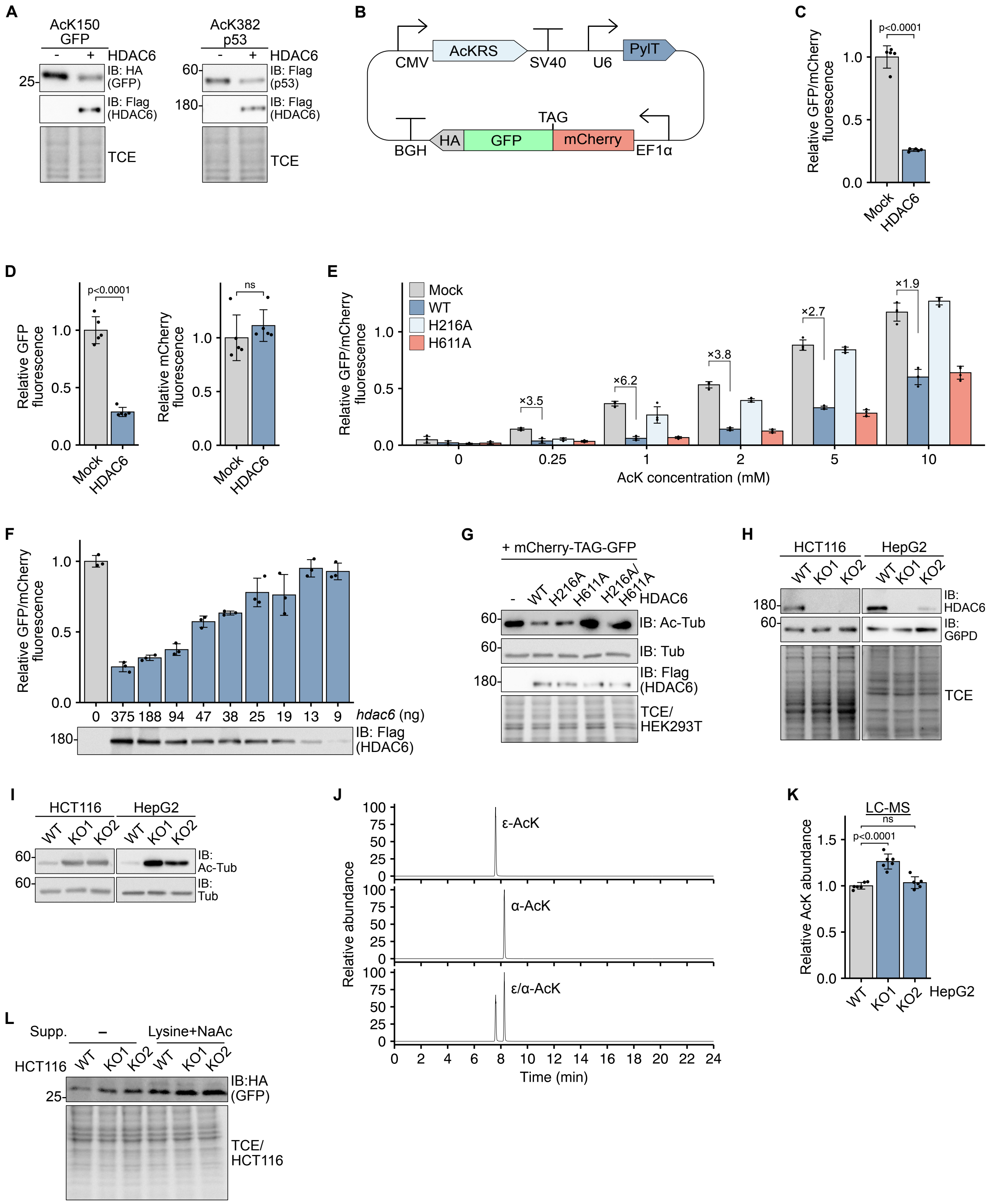
A
AcK150 GFP
AcK382 p53
HDAC6
IB: HA (GFP)
IB: Flag (HDAC6)
IB: Flag (p53)
TCE
B
CMV
AcKRS
SV40
U6
PylT
TAG
BGH
HA
GFP
mCherry
EF1α
C
p<0.0001
Relative GFP/mCherry fluorescence
Mock
HDAC6
D
Relative GFP fluorescence
Relative mCherry fluorescence
ns
E
Mock
WT
H216A
H611A
×3.5
×6.2
×3.8
×2.7
×1.9
AcK concentration (mM)
F
hdac6 (ng)
IB: Flag (HDAC6)
G
+ mCherry-TAG-GFP
IB: Ac-Tub
IB: Tub
TCE/ HEK293T
H
HCT116
HepG2
WT
KO1
KO2
IB: HDAC6
IB: G6PD
I
IB: Ac-Tub
IB: Tub
J
ε-AcK
α-AcK
ε/α-AcK
Relative abundance
Time (min)
K
LC-MS
Relative AcK abundance
L
Supp.
Lysine+NaAc
IB:HA (GFP)
TCE/ HCT116

**Figure S2**

**Figure S2. Additional characterization of HDAC6 CD1 activity in cells, related to Figure 1.**
(A) Immunoblotting using anti-Flag antibody showing the levels of full-length C-terminal HA-tagged GFP or Flag-tagged p53 with an in-frame TAG mutation at position 150 or 382, respectively, coexpressed with HDAC6-Flag or expressed without HDAC6 (empty vector). The efficiency of TAG-encoded AcK incorporation, and consequently, expression of full-length proteins, is dependent on AcK availability.
(B) Schematic representation of a single-plasmid-based expression system for a dual-color fluorescence assay to evaluate AcK incorporation efficiency in mammalian cells. AcKRS, PylT, and mCherry-TAG-GFP are expressed under the control of a CMV promoter, U6 promoter, and EF1α promoter, respectively.
(C) Relative GFP/mCherry fluorescence ratio measured in clear lysates of HEK293T cells cultured with 2 mM AcK and cotransfected with mCherry-TAG-GFP encoding plasmid and HDAC6-Flag encoding plasmid or empty vector.
(D) Relative GFP (left) or mCherry (right) fluorescence measured in cleared lysates of HEK293T cells cultured with 2 mM AcK and cotransfected with mCherry-TAG-GFP encoding plasmid and HDAC6-Flag encoding plasmid or empty vector.
(E) Relative GFP/mCherry fluorescence ratio measured in clear lysates of HEK293T cells cultured with increasing AcK concentrations and cotransfected with mCherry-TAG-GFP encoding plasmid and WT or mutant HDAC6-Flag encoding plasmid or empty vector.
(F) Relative GFP/mCherry fluorescence ratio measured in clear lysates of HEK293T cells cultured with 2 mM AcK and cotransfected with 375 ng mCherry-TAG-GFP encoding plasmid and indicated amount of WT HDAC6-Flag encoding plasmid. The total amount of transfected DNA was kept constant (750 ng) using an empty vector.
(G) Immunoblotting using the indicated antibodies, showing levels of K40-acetylated α-tubulin in cleared lysates of HEK293T cells cotransfected with the mCherry-TAG-GFP encoding plasmid and WT or mutant HDAC6-Flag encoding plasmid or empty vector.
(H) Immunoblotting using anti-HDAC6 and anti-G6PD antibodies showing the expression levels of HDAC6 in WT and HDAC6-knockout HCT116 and HepG2 cell lines. KO1 and KO2 cell lines were created using sg1 and sg2 guide RNA sequences (as detailed in the Materials and Methods section), respectively.
(I) Immunoblotting using anti-K40-acetylated α-tubulin and anti-α-tubulin antibodies, showing the level of α-tubulin K40 acetylation in WT versus HDAC6-knockout HCT116 and HepG2 cell lines.
(J) LC-MS analysis showing the difference in elution time between $N^{\alpha}$- and $N^{\varepsilon}$-acetyl-L-lysine standards. The clear separation between the two structural isomers enables the quantification of $N^{\varepsilon}$-acetyllysine abundance in cells.
(K) Relative abundance of endogenous AcK in WT or HDAC6 knockout HepG2 cell lines, quantified by LC-MS.
(L) Immunoblotting using anti-HA antibody showing the expression levels of full-length GFP-HA in WT and HDAC6-knockout HCT116 cell lines, transfected with GFP-N150TAG-HA and AcKRS encoding plasmid. Cells were treated with 8 mM lysine and 5 mM NaAc and cultured without supplemented AcK. Full-length GFP expression levels report on the abundance of endogenous AcK.
Data represent the mean ± S.D. of at least three independent experiments. Statistical analysis was performed using ANOVA followed by Tukey post hoc analysis (K) or using the Welch two-sample t-test (C, D). ns, non-significant. In-gel TCE fluorescence serves as a loading control.

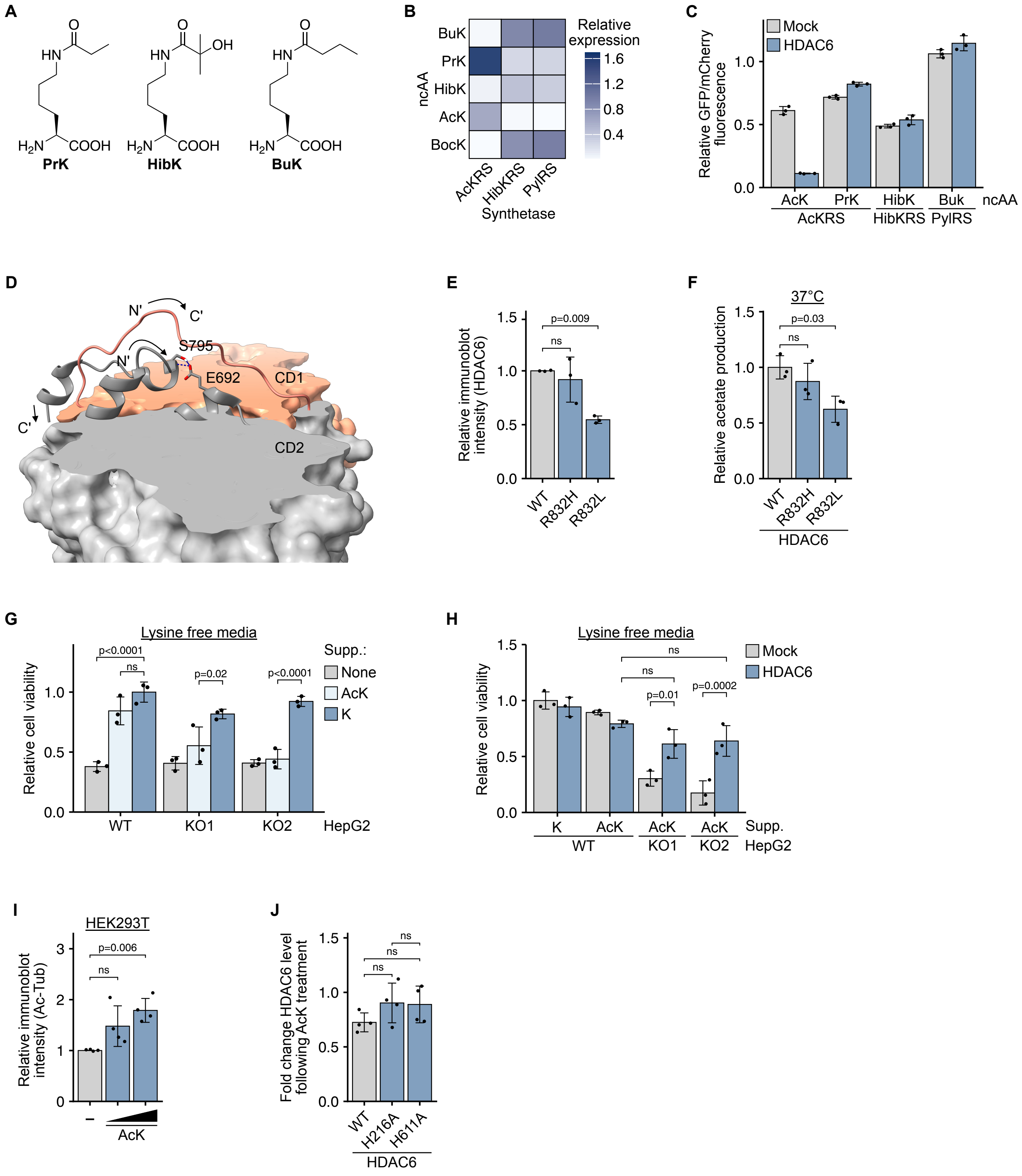
A
PrK
HibK
BuK
B
Relative expression
ncAA
Synthetase
AcKRS
HibKRS
PylRS
C
Mock
HDAC6
Relative GFP/mCherry fluorescence
D
S795
E692
CD1
CD2
E
Relative immunoblot intensity (HDAC6)
F
37°C
Relative acetate production
G
Lysine free media
Relative cell viability
Supp.:
None
AcK
K
HepG2
H
Lysine free media
I
HEK293T
Relative immunoblot intensity (Ac-Tub)
J
Fold change HDAC6 level following AcK treatment

**Figure S3**

**Figure S3. Additional characterization of AcK physiological significance, related to Figure 2.**
(A) Chemical structures of propionyl lysine (PrK), 2-hydroxyisobutyryl lysine (HibK), and butyryl lysine (BuK).
(B) Evaluation of acyl-lysine incorporation efficiency by WT PylRS, or synthetases evolved for the incorporation of AcK (AcKRS) or HibK (HibKRS). Cells were cultured with the indicated acyl-lysine (2 mM) or Nε tert-butyloxycarbonyl protected lysine (BocK; 1 mM) and cotransfected with mCherry-TAG-GFP and indicated synthetase encoding plasmids. Incorporation efficiency was evaluated by measuring the GFP/mCherry fluorescence ratio, and indicated by the color scheme.
(C) Relative GFP/mCherry fluorescence ratio in lysates of HEK293T cells cultured with the indicated acyl-lysine and cotransfected with mCherry-TAG-GFP encoding plasmid, the suitable synthetase encoding plasmid (determined according to panel B), and HDAC6-Flag or empty vector.
(D) Crystal structure of *D. rerio* HDAC6 (PDB ID: 7QNO).[2] CD1 and CD2 are colored in orange and grey, respectively. Parts of the structure were removed (indicated by the smooth surface), to expose the interaction between S795 and E692 (R832 and E729 in human HDAC6).
(E) Relative HDAC6-Flag levels in HEK293T cells transfected with a plasmid encoding the expression of WT or natural HDAC6-Flag variants. Bars represent the relative HDAC6-Flag immunoblot intensity.
(F) Relative acetate production following 1 hr incubation at 37°C, of 3 mM AcK with cleared lysates of HEK293T cells transfected with WT or mutant HDAC6-Flag expressing plasmid.
(G) Relative viability of WT and HDAC6-knockout HepG2 cells cultured in lysine-deficient media supplemented with 0.8 mM lysine or AcK, or without supplements. Data are presented relative to WT cells cultured with lysine.
(H) Relative viability of WT and HDAC6-knockout HepG2 cells cultured in lysine-free media, as a function of lysine or AcK supplement (as described in G), and HDAC6 expression. Data are presented relative to mock-transfected WT cells cultured with lysine.
(I) Relative α-tubulin K40 acetylation levels in HEK293T cells as a function of AcK concentration (0, 2, and 5 mM). Bars represent the ratio between AcK40 α-tubulin and α-tubulin immunoblot intensities.
(J) Fold change in HDAC6-Flag levels in HDAC6-knockout HCT116 cells transfected with WT or mutant HDAC6-Flag, in response to 5 mM AcK treatment. Quantified by immunoblotting.
Data represent the mean ± S.D. of at least three independent experiments. Statistical analysis was performed using ANOVA followed by Tukey post hoc analysis (E–J). ns, non-significant.

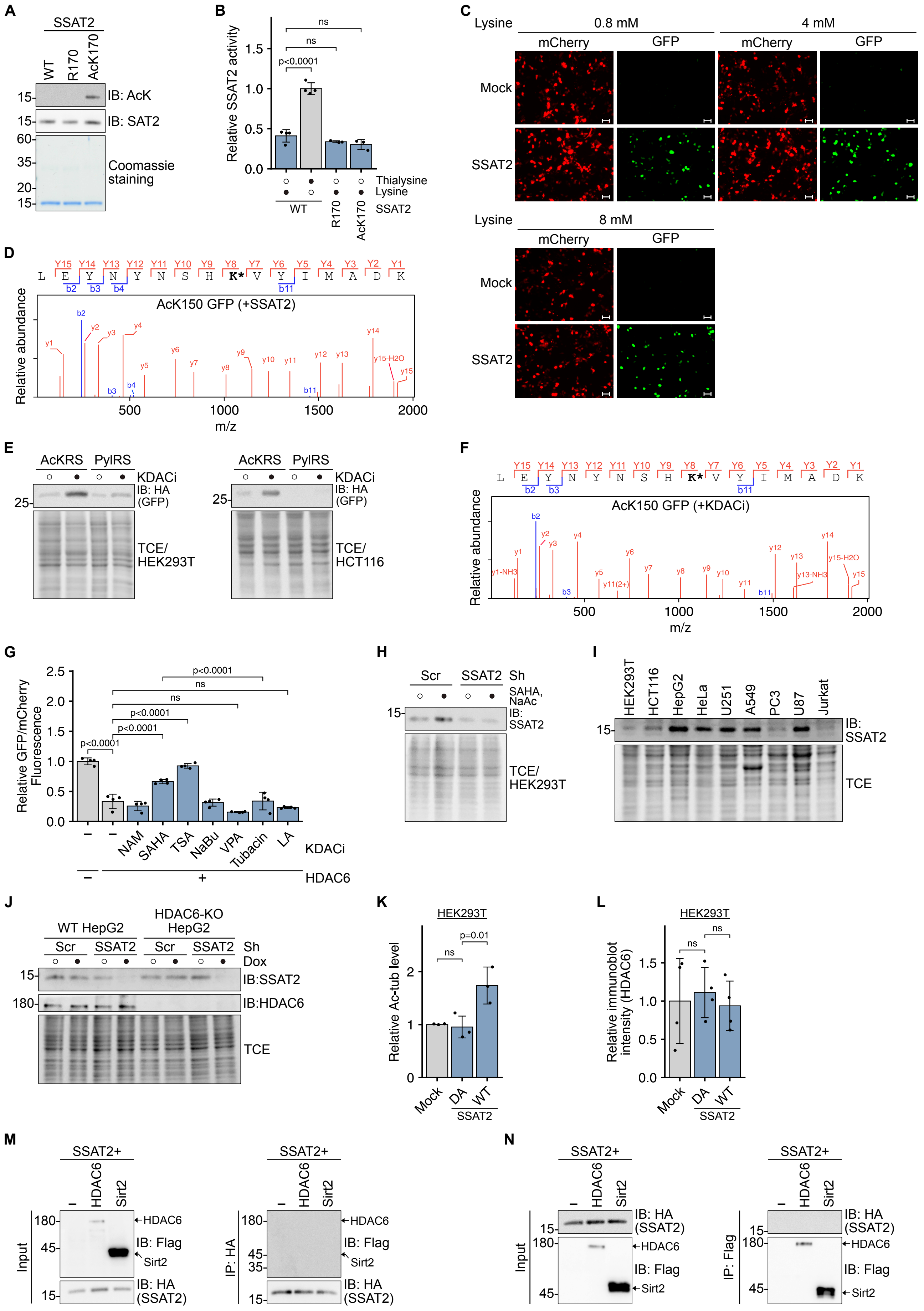
A
SSAT2
WT
R170
AcK170
IB: AcK
IB: SAT2
Coomassie staining
B
Relative SSAT2 activity
Thialysine
Lysine
SSAT2
WT
R170
AcK170
p<0.0001
ns
C
Lysine
0.8 mM
4 mM
8 mM
mCherry
GFP
Mock
SSAT2
D
AcK150 GFP (+SSAT2)
Relative abundance
m/z
E
AcKRS
PylRS
KDACi
IB: HA (GFP)
TCE/ HEK293T
TCE/ HCT116
F
AcK150 GFP (+KDACi)
Relative abundance
m/z
G
Relative GFP/mCherry Fluorescence
NAM
SAHA
TSA
NaBu
VPA
Tubacin
LA
KDACi
HDAC6
H
Scr
SSAT2
Sh
SAHA, NaAc
IB: SSAT2
TCE/ HEK293T
I
HEK293T
HCT116
HepG2
HeLa
U251
A549
PC3
U87
Jurkat
IB: SSAT2
TCE
J
WT HepG2
HDAC6-KO HepG2
Scr
SSAT2
Sh
Dox
IB:SSAT2
IB:HDAC6
TCE
K
HEK293T
Relative Ac-tub level
p=0.01
ns
Mock
DA
WT
SSAT2
L
HEK293T
Relative immunoblot intensity (HDAC6)
ns
Mock
DA
WT
SSAT2
M
SSAT2+
HDAC6
Sirt2
Input
IP: HA
IB: Flag
IB: HA (SSAT2)
N
SSAT2+
Input
IP: Flag
IB: HA (SSAT2)
IB: Flag

Figure S4

**Figure S4. Additional characterization of cellular AcK synthesis, related to Figure 3.**
(A) Immunoblotting using anti-AcK and anti-SSAT2 antibodies, and Coomassie blue staining of purified WT and SSAT2 variants expressed in *E. coli.*
(B) Relative in vitro acetyltransferase activity of purified recombinant SSAT2 (WT, R170, or AcK170) incubated for 1 hr with (•) or without (○) the indicated substrate. Bars represent the amount of released CoA quantified using DTNB, normalized to the value measured for WT SSAT2 incubated with thialysine.
(C) Fluorescence imaging of HEK293T cells cotransfected with a plasmid encoding the expression mCherry-TAG-GFP, and HA-SSAT2 or empty vector, and cultured in the presence of 0.8, 4, or 8 mM lysine. Scale bar, 20 µm.
(D) LC-MS/MS analysis of immunopurified AcK150GFP-HA expressed as described in Figure 3A. The sequence of the identified tryptic peptide shows the position of incorporated AcK (indicated by **K***). The sequences and masses of detected b and y fragments are presented.
(E) Immunoblotting using anti-HA antibody showing the expression level of full-length GFP-HA in HEK293T and HCT116 cells as a function of KDACi (SAHA and NAM) treatment and coexpressed synthetase. Expression is enabled by the incorporation of endogenous AcK into GFP-150TAG-HA. The lack of AcK incorporation by PylRS, demonstrates that KDACi did not affect the fidelity of the amber-suppression system or promoted readthrough of the TAG stop codon at position 150.
(F) LC-MS/MS analysis of immunopurified AcK150GFP-HA expressed as described in E. The sequence of the identified tryptic peptide shows the position of incorporated AcK (indicated by **K***). The sequences and masses of detected b and y fragments are presented.
(G) Relative GFP/mCherry fluorescence ratio in lysates of HEK293T cells cultured with the indicated compounds and cotransfected with mCherry-TAG-GFP encoding plasmid, and HDAC6-Flag or empty vector. TSA: trichostatin A; NaBu: sodium butyrate; VPA: valproic acid sodium salt; LA: R-(+)-alpha-lipoic acid.
(H) Immunoblotting using anti-SSAT2 antibody, showing the level of endogenous SSAT2 in HEK293T with stable Dox-dependent expression of Sh against SSAT2 or scrambled Sh sequence. Cells were cultured in the presence (•) or absence (○) of SAHA and NaAc.
(I) Immunoblotting using anti-SSAT2 antibody showing the level of endogenous SSAT2 in the indicated cell lines.
(J) Immunoblotting using anti-SSAT2 and anti-HDAC6 antibodies showing the levels of endogenous SSAT2 and HDAC6 in WT or HDAC6-knockout HepG2 cells with stable Dox-dependent expression of Sh against SSAT2 or scrambled Sh sequence.
(K) Relative α-tubulin K40 acetylation levels in HEK293T cells transfected with WT or inactive (DA) HA-SSAT2 expressing plasmid, or empty vector. Bars represent the ratio between AcK40 α-tubulin and α-tubulin immunoblot intensities.
(L) Immunoblotting using anti-HDAC6 antibody showing the level of endogenous HDAC6 in HEK293T cells transfected with WT or mutant HA-SSAT2 expressing plasmid or empty vector.
(M, N) Reciprocal co-IP to detect a stable interaction between HA-SSAT2 and HDAC6-Flag or Sirt2-Flag. Immunoprecipitation from cleared lysates of HEK293T cells cotransfected with HA-SSAT2 and indicated KDAC expressing plasmids was performed using either an HA antibody against HA-SSAT2 (M) or a Flag antibody against HDAC6-Flag or Sirt2-Flag (N).
Data represent the mean ± S.D. of at least three independent experiments. Statistical analysis was performed using ANOVA followed by Tukey post hoc analysis (B, G, K, L). ns, non-significant. Scr, scrambled Sh sequence. In-gel TCE fluorescence serves as a loading control.

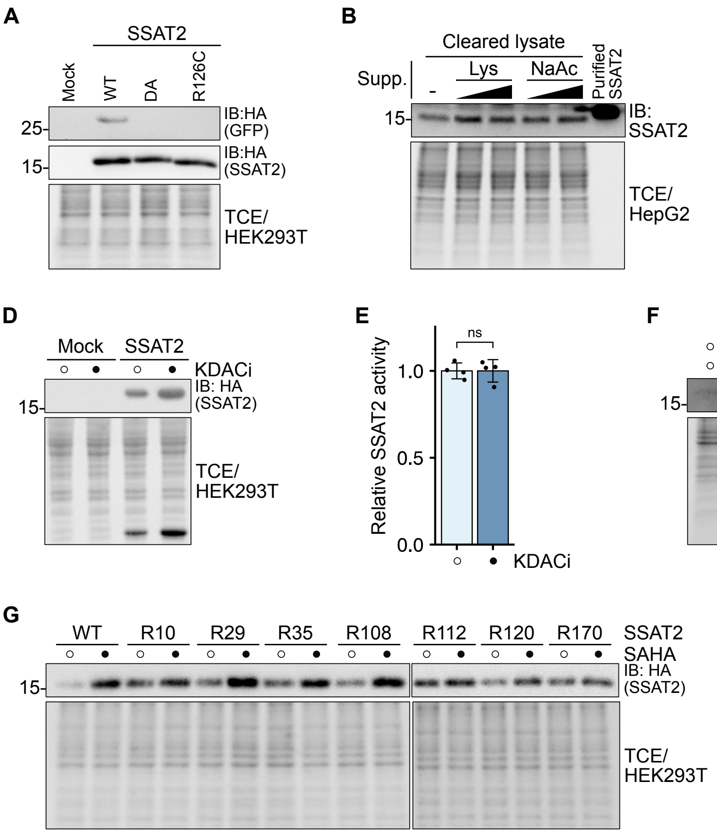


**Figure S5. Additional characterization of SSAT2 acetylation, related to Figure 3.**
(A) Immunoblotting using anti-HA antibody showing the level of full-length GFP-HA in cells cotransfected with plasmid encoding GFP-N150TAG-HA and AcKRS and plasmid encoding the indicated HA-SSAT2 variant or empty vector.
(B) Immunoblotting using anti-SSAT2 antibody showing the levels of endogenous SSAT2 in HepG2 cells cultured in normal media without additional supplements, or media supplemented with increasing concentrations of lysine (4 mM and 8 mM) or NaAc (5 mM and 10 mM). Purified recombinant 6×His-SSAT2 was used as a positive control.
(C) Immunoblotting using anti-GFP antibody showing the level of GFP expressed in HepG2 cells cultured in normal media without additional supplements, or media supplemented with increasing concentrations of lysine (4 mM and 8 mM) or NaAc (5 mM and 10 mM). GFP expression levels were not affected by the addition of lysine or NaAc.
(D) Immunoblotting using anti-HA antibody showing the levels of HA-SSAT2 in HEK293T cells transfected with HA-SSAT2 expressing plasmid or empty vector and cultured with (•) or without (○) KDACi (NAM and SAHA). Exogenous HA-SSAT2 is stabilized by KDACi.
(E) Relative in vitro acetyltransferase activity of Flag-isolated SSAT2 expressed in HEK293T cells cultured with or without KDACi (NAM and SAHA) for 2 days. SSAT2 activity is not affected by KDACi.
(F) Immunoblotting using anti-SSAT2 antibody showing the stabilizing effect of SAHA on levels of endogenous SSAT2 in both HEK293T and HepG2 cell lines.
(G) Immunoblotting using anti-HA antibody showing the stabilizing effect of SAHA on HA-SSAT2 levels in HEK293T cells transfected with a plasmid encoding the expression of WT or mutant HA-SSAT2.
Data represent the mean ± S.D. of at least three independent experiments. Statistical analysis was performed using the Welch two-sample t-test (E). ns, non-significant. In-gel TCE fluorescence serves as a loading control.

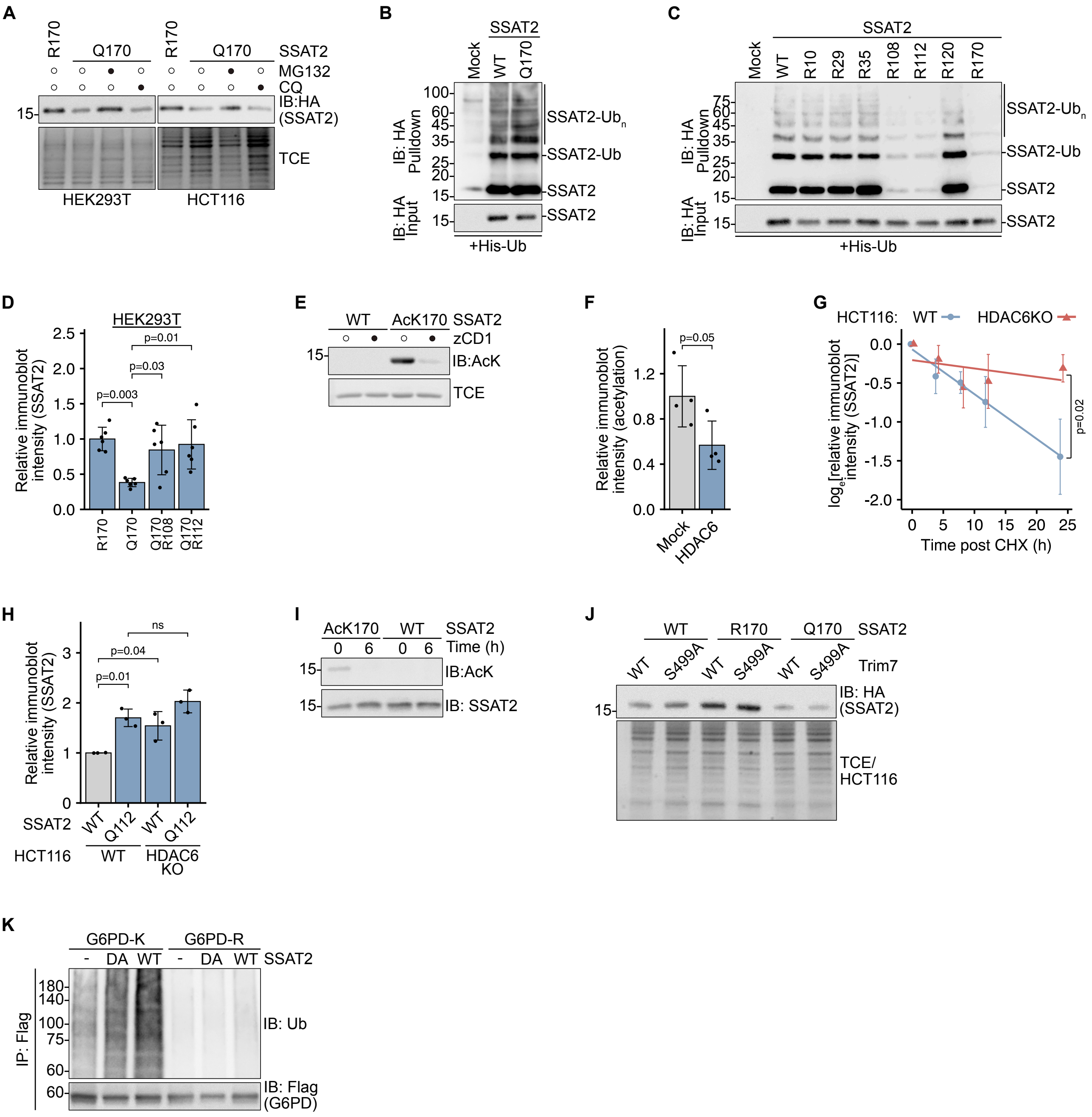
A
R170
Q170
R170
Q170
SSAT2
MG132
CQ
IB:HA (SSAT2)
TCE
HEK293T
HCT116
B
SSAT2
Mock
WT
Q170
IB: HA Pulldown
IB: HA Input
SSAT2-Ubn
SSAT2-Ub
SSAT2
+His-Ub
C
Mock
WT
R10
R29
R35
R108
R112
R120
R170
D
HEK293T
Relative immunoblot intensity (SSAT2)
p=0.003
p=0.03
p=0.01
E
WT
AcK170
zCD1
IB:AcK
F
Relative immunoblot intensity (acetylation)
p=0.05
Mock
HDAC6
G
HCT116: WT HDAC6KO
loge[relative immunoblot intensity (SSAT2)]
Time post CHX (h)
p=0.02
H
p=0.01
p=0.04
ns
Q112
HCT116
HDAC6 KO
I
Time (h)
IB: SSAT2
J
Trim7
S499A
IB: HA (SSAT2)
TCE/ HCT116
K
G6PD-K
G6PD-R
DA
IP: Flag
IB: Ub
IB: Flag (G6PD)

**Figure S6**

**Figure S6. Additional characterization of SSAT2 function, related to Figure 4.**
(A) Immunoblotting using anti-HA antibody showing the levels of R170 or Q170 HA-SSAT2 mutants expressed in HEK293T or HCT116 cells treated with 5 µM MG132 or 50 µM chloroquine for 16 h before harvest.
(B) Immunoblotting using anti-HA antibody showing the levels of WT and Q170 HA-SSAT2 in proteins isolated, using Ni-NTA resin, from cleared lysates of HEK293T cells cotransfected with plasmid encoding HA-SSAT2 and 6×His-Ub or empty vector.
(C) Immunoblotting using anti-HA antibody showing the levels of indicated HA-SSAT2 variants in proteins isolated, using cobalt resin, from lysates of HEK293T cells cotransfected with plasmid encoding HA-SSAT2 and 6×His-Ub or empty vector.
(D) Levels of indicated HA-SSAT2 mutants expressed in HEK293T cells. Bars represent the relative HA-SSAT2 immunoblot intensities.
(E) Immunoblotting using anti-AcK antibody showing the acetylation levels of purified WT and AcK170 SSAT2 expressed in *E. coli*, following incubation with (•) or without (◦) zCD1.
(F) Acetylation levels of purified AcK112 SSAT2 expressed in *E. coli*, following incubation with cleared lysates of HEK293T cells transfected with HDAC6-Flag expressing plasmid or empty vector. Bars represent the relative K112-acetylated SSAT2 immunoblot intensities.
(G) Log relative HA-SSAT2 levels expressed in WT and HDAC6-knockout HCT116, described as a function of time post-CHX treatment. Data are reported as mean ± S.E.M.
(H) Levels of WT and Q112 HA-SSAT2 expressed in WT or HDAC6-knockout HCT116 cells. Bars represent the relative HA-SSAT2 immunoblot intensities.
(I) Immunoblotting using anti-AcK antibody showing the acetylation levels of WT and AcK170 SSAT2 before and 6 h after delivery into WT HCT cells by electroporation.
(J) Immunoblotting using anti-HA antibody showing the levels of indicated HA-SSAT2 variant coexpressed in HCT116 cells with WT Trim7, or the inactive S499A Trim7 mutant.
(K) Immunoblotting using anti-ubiquitin antibody showing the poly-ubiquitylation of indicated G6PD-Flag variants in IPed proteins isolated, using anti-Flag antibody, from lysates of HEK293T cells cotransfected with plasmid encoding G6PD-Flag and 6×His-Ub or empty vector.
Data represent the mean ± S.D. of at least three independent experiments, unless indicated otherwise. Statistical analysis was performed using the Welch two-sample t-test (F), ANOVA followed by Tukey post hoc analysis (D, H), or pairwise trend comparisons with $p$-values adjusted for multiple comparisons using the Holm method (G). In-gel TCE fluorescence serves as a loading control.

### Supplementary References


1. Yin, X., Chan, L.S., Bose, D., Jackson, A.U., VandeHaar, P., Locke, A.E., Fuchsberger, C., Stringham, H.M., Welch, R., Yu, K., Fernandes Silva, L., Service, S.K., Zhang, D., Hector, E.C., Young, E., Ganel, L., Das, I., Abel, H., Erdos, M.R., Bonnycastle, L.L., Kuusisto, J., Stitziel, N.O., Hall, I.M., Wagner, G.R., FinnGen, Kang, J., Morrison, J., Burant, C.F., Collins, F.S., Ripatti, S., Palotie, A., Freimer, N.B., Mohlke, K.L., Scott, L.J., Wen, X., Fauman, E.B., Laakso, M., and Boehnke, M. (2022). Genome-Wide Association Studies of Metabolites in Finnish Men Identify Disease-Relevant Loci. Nature Communications *13*, 1644. http://doi.org/10.1038/s41467-022-29143-5.

2. Langousis, G., Sanchez, J., Kempf, G., and Matthias, P. (2023). Expression and Crystallization of HDAC6 Tandem Catalytic Domains. Methods in Molecular Biology (clifton, N.J.) *2589*, 467–480. http://doi.org/10.1007/978-1-0716-2788-4_30.
